# The gamma-Protocadherins regulate the survival of GABAergic interneurons during developmentally-regulated cell death

**DOI:** 10.1101/2020.01.15.908087

**Authors:** Candace H. Carriere, Anson D. Sing, Wendy Xueyi Wang, Brian E. Jones, Yohan Yee, Jacob Ellegood, Julie Marocha, Harinad Maganti, Lola Awofala, Amar Aziz, Jason P. Lerch, Julie L. Lefebvre

## Abstract

Inhibitory interneurons integrate into developing circuits in specific ratios and distributions. In the cortex, the formation of inhibitory networks occurs concurrently with the apoptotic elimination of a third of GABAergic interneurons. The molecular mechanisms that select GABAergic interneurons to survive or die are unknown. Here we report that the clustered Protocadherins regulate GABAergic cell survival in the developing brain. Deletion of the *Pcdh-gamma* genes (Pcdhgs) from GABAergic neurons in mice causes a severe loss of inhibitory neurons in multiple brain regions and results in motor deficits and seizure activities. By focusing on the neocortex and cerebellar cortex, we demonstrate that GABAergic interneuron loss results from elevated apoptosis during the postnatal wave of *Bax-*dependent programmed cell death. Pro-survival AKT signals are reduced in *Pcdhg-*deficient interneurons, diminishing the intrinsic capacity of interneurons to compete and incorporate into developing networks. We propose that the Pcdhgs mediate selective GABAergic interneuron survival to contribute to the formation of balanced inhibitory networks.

## INTRODUCTION

The optimal function of neuronal networks is dependent on balanced populations of excitatory and inhibitory neurons. Local GABAergic-releasing interneurons are integrated at particular ratios to match the principal neuronal populations and their circuit requirements. Establishing appropriate interneuron densities is critical to circuit maturation and function, as the specialized forms of synaptic inhibition conferred by these cells shape principal neuron firing and network synchronization (Le Magueresse and Monyer, 2013). Alterations in GABAergic interneuron density and maturation are implicated in several human neurodevelopmental disorders and mouse models (Donovan and Basson, 2017; Edamakanti et al., 2018; Fu et al., 2012; Kataoka et al., 2010; Lacaille et al., 2019; Marin, 2012; Penagarikano et al., 2011)

Programmed cell death (PCD) is critical for adjusting neuronal population sizes and eliminating improperly connected cells in the developing brain (Wong and Marin, 2019). However, the extent to which inhibitory interneurons are sculpted by PCD was demonstrated only recently. In mice, 30-40% of prospective neocortical GABAergic interneurons are eliminated between postnatal days 5 and 10 (Southwell et al., 2012). Cortical interneuron (cIN) apoptosis occurs through the Bax-dependent intrinsic pathway and affects the major classes deriving from the medial and caudal ganglionic eminences (MGE and CGE, respectively) (Denaxa et al., 2018; Priya et al., 2018; Southwell et al., 2012; Wong et al., 2018). cIN PCD occurs on the heels of pyramidal PCD which tapers by P5 (Blanquie et al., 2017; Miller, 1995; Verney et al., 2000; Wong et al., 2018). Matching between cINs and pyramidal neurons might be determined by competition for target-derived neurotrophic factors, as in the peripheral nervous system (Dekkers et al., 2013; Deppmann et al., 2008; Snider, 1994). However, cIN number is not affected by manipulations of TrkB or BDNF (Priya et al., 2018; Sanchez-Huertas and Rico, 2011; Southwell et al., 2012), suggesting the regulation of cIN survival by other mechanisms. Neural activity is one critical factor that influences the extent of inhibitory and excitatory PCD in the cortex by modulating pro-survival pathways such as PI3K/Akt and calcineurin/NFAT (Hardingham et al., 2002; Priya et al., 2018; Wagner-Golbs and Luhmann, 2012; Wong et al., 2018), and suppressing pro-apoptotic effectors (Golbs et al., 2011; Heck et al., 2008; Leveille et al., 2010). Blocking cIN activity by NMDA antagonists or chemogenetic approaches accentuates cIN apoptotic loss while depolarization increases the number of surviving cells (Denaxa et al., 2018; Priya et al., 2018; Wang et al., 2017). Moreover, increasing the activity of pyramidal neurons or simply their numbers are sufficient to expand the cIN population (Wong et al., 2018). However, the influence of activity is cIN class-specific (Priya et al., 2018), and cell- or population-intrinsic factors also contribute to cIN survival (Southwell et al., 2012). Thus, life-or-death decisions among developing GABAergic interneurons are regulated by multiple selective interactions but the molecules that mediate them are unknown.

Elsewhere in the CNS, the clustered Protocadherins (cPcdhs) are essential regulators for interneuronal survival during PCD in the spinal cord and retina (Hasegawa et al., 2016; Ing-Esteves et al., 2018; Lefebvre et al., 2008; Prasad et al., 2008; Wang et al., 2002; Weiner et al., 2005). In mice, the cPcdhs comprise a large family of 58 cadherin-related transmembrane molecules that are encoded by three tandemly-arranged gene clusters, *Pcdh-alpha (Pcdha), Pcdh-beta (Pcdhb)*, and *Pcdh-gamma (Pcdhg)* (Wu and Maniatis, 1999). The cPcdhs are noted for their potential for immense cell surface diversity and selectivity to mediate complex neuronal interactions (Lefebvre, 2017; Mountoufaris et al., 2018). Consistent with this idea, cPcdhs play diverse roles in neurite patterning such as arborization, tiling, and self-avoidance which are proposed to result from homophilic cPcdh interactions that mediate adhesion, or self/non-self discrimination and repulsion (Chen et al., 2017; Garrett et al., 2012; Ing-Esteves et al., 2018; Katori et al., 2009; Lefebvre et al., 2012; Molumby et al., 2016; Mountoufaris et al., 2017). The cPcdhs also regulate neuronal survival in a genetically distinct pathway. *Pcdh-gamma* and triple *cPcdh* knockout mice die at birth due to massive apoptosis in the hindbrain and spinal cord (Hasegawa et al., 2016; Wang et al., 2002). Conditional deletions in the retina and spinal cord revealed that Pcdhgs mediate a survival program that follows the developmental pattern of Bax-dependent PCD (Lefebvre et al., 2008; Prasad et al., 2008). Of the twenty-two *Pcdhgs*, the three C-type Pcdhg genes are required for neuronal survival (Chen et al., 2012) and a recent study of CRISPR-mediated *Pcdhg* deletion mutants suggests that PcdhgC4 is solely essential (Garrett et al., 2019). Constitutive *Pcdha* and *Pcdhb* mutants, on the other hand, are viable and exhibit little to no cell loss (Hasegawa et al., 2016; Ing-Esteves et al., 2018). But neuronal death phenotypes are profoundly exacerbated in double *Pcdha; Pcdhg* mutant retinas (Ing-Esteves et al., 2018) and triple *cPcdh* knockout spinal cords (Hasegawa et al., 2016), revealing complex redundancy and cooperation between the cPcdhs in survival.

Here we test the hypothesis that the Pcdhgs regulate the survival of GABAergic interneuron populations during PCD in the developing brain. cPcdhs are dispensable the survival of major principal neuron populations including cortical pyramidal neurons, GABAergic cerebellar Purkinje cells, and serotonergic projection neurons (Chen et al., 2017; Garrett et al., 2012; Katori et al., 2009; Lefebvre et al., 2012). Using the pan-GABAergic Gad2-Cre driver (Taniguchi et al., 2011) to broadly delete the *Pcdhgs* from inhibitory neuronal populations, we find that GABAergic populations are profoundly reduced in size in multiple brain regions. Consequently, GABAergic *Pcdhg* mutant mice develop behavioral deficits and seizures. Focusing on interneuron populations in the cortex and cerebellum, we demonstrate that the Pcdhgs mediate a pro-survival mechanism for the specific control of interneuron survival during developmentally-regulated cell death.

## RESULTS

### GABAergic-specific deletion of *Pcdhgs* in mice causes neurological and behavioral deficits

To investigate the role of the Pcdhgs in inhibitory neuron development, we used a pan-GABAergic Cre line to delete *Pcdhg* from broad classes of differentiating GABAergic cells. Conditional *Pcdhg^f^* mice (Lefebvre et al., 2008; Prasad et al., 2008) were crossed to Gad2-ires-Cre (Taniguchi, 2014; Taniguchi et al., 2011) which is active in the brain by late embryonic stages (Saito et al., 2019). Homozygous *Pcdhg^f/f^*; *Gad2-ires-Cre* mutant offspring (herein referred to as *Pcdhg^GcKO^*) were viable at P0 but displayed pronounced growth and neurological deficits within the first few postnatal weeks. By postnatal day 28, *Pcdhg^GcKO^* mutants were on average 30% smaller in body weight than control littermates (Figure 1A; 16.6g compared to 10.8, N = 15 animals per genotype, *p* = 0.00009, *t*-test). Juvenile *Pcdhg^GcKO^* mutant mice exhibited forelimb and hindlimb clasping upon tail suspension, a motor impairment phenotype that occurred with 100% penetrance by P21 and persisted at later ages (Figure 1B). *Pcdhg^GcKO^* motor impairments were also indicated by reduced performance on the accelerating rotarod (Figure 1C).

**Figure 1.**
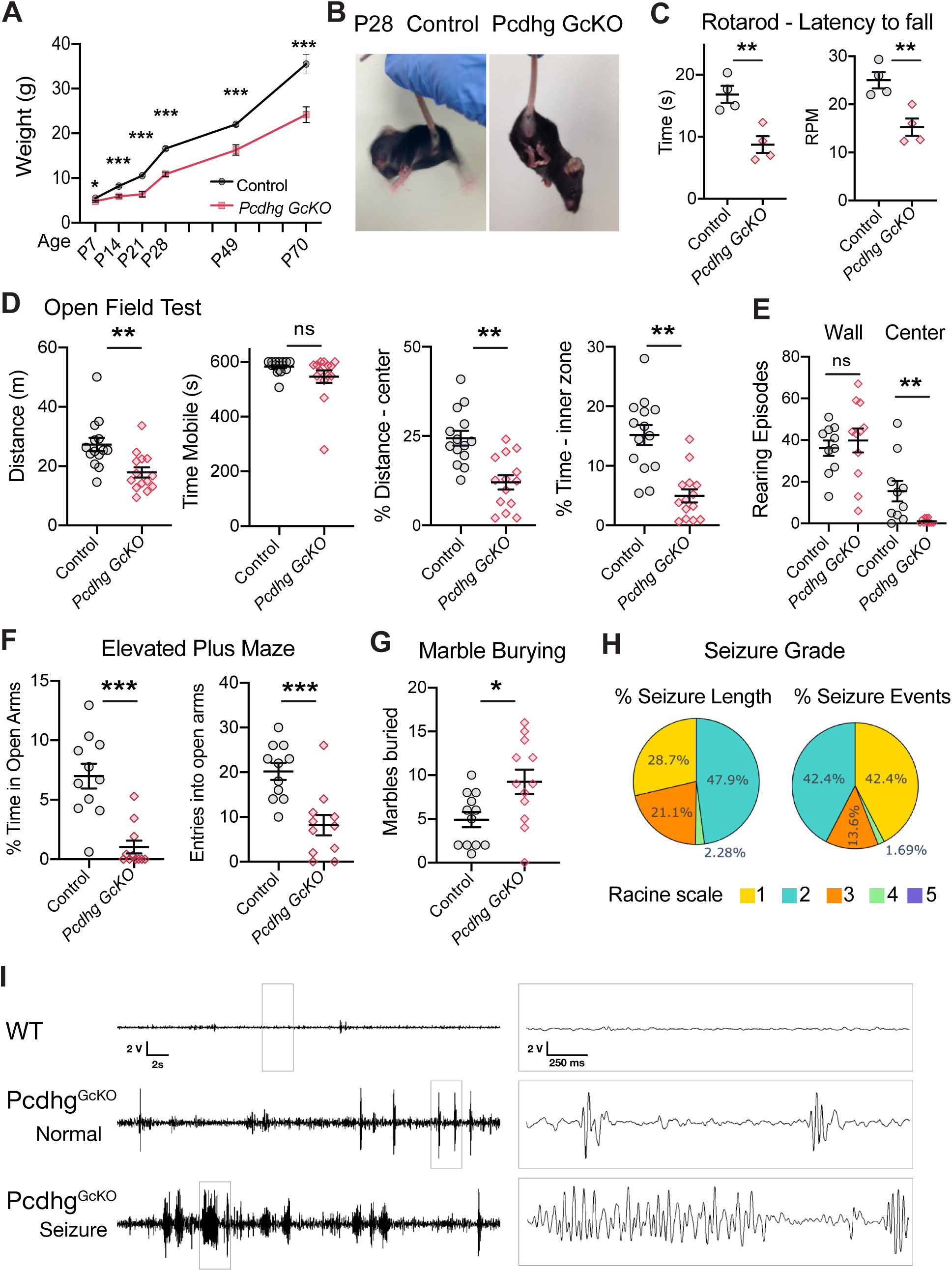
GABAergic *Pcdhg* mutants are smaller and exhibit neurological and behavioral deficits. **A.** Weights of control (black line) and *Pcdhg GcKO* (red line) mice. **B.** *Left*, P28 WT sibling. *Right, Pcdhg GcKO* mutant displays forelimb and hindlimb clasping. **C-G.** Assays for motor deficits and anxiety-like behaviors of control (black) and *Pcdhg GcKO* mice (red). Data are presented as scatter plots with distribution of test animals, and whisker plots denoting mean with SEM. **C.** Rotarod. *Pcdhg GcKO* mice exhibit decreased time to fall (*left*, *p=*0.0057) and rotarod speed (*right*, *p =* 0.0074, Student’s unpaired t test. N = 4 animals per genotype). **D.** Open Field Test. *Pcdhg GcKO* mice exhibit reduced total distance traveled (*left*, *p=*0.001), similar moving time (ns, *p* =0.10), reduced percent distance traveled (*p =*0.00027), and percent time spent in the inner zone (*right*, *p =* 0.00005, Mann-Whitney U-test, N = 14 animals per genotype). **E.** Rearing. *Pcdhg GcKO* perform similar wall climbs as control (*p* = 0.31) but reduced rears in the open arena (43.20 mean episodes in control vs 1.2 in *Pcdhg GcKO, p =* 0.0017, Mann-Whitney U-test; N = 10 animals per genotype). **F.** Elevated Plus Maze. *Pcdhg GcKO* mice spend less time (*p =* 0.00008) and make fewer entries into open arms (*p =* 0.0008, Mann-Whitney U-test; N = 11 animals per genotype). **G.** Numbers of marbles buries were greater for *Pcdhg GcKO* than control mice (*p =* 0.022, Mann-Whitney U-test; N =12 animals per group). **H.** Assessment of total seizure time and events of *Pcdhg GcKO* mice (N = 4) by Racine scale. **I.** Examples of EEG waveforms of cortical baseline activities recorded with bipolar electrodes in WT and *Pcdhg GcKO* mice (N = 4 per genotype). * *p* < 0.05; ** *p* < 0.01; *** *p* < 0.001.

We analyzed the locomotor and exploratory behaviors of GABAergic *Pcdhg^GcKO^* mutant mice (2-3 months of age) in open field and elevated plus maze tests. *Pcdhg^GcKO^* mutants traveled shorter distances but were not generally hypoactive, as they did not differ significantly from control in total moving time and wall climbs (Figure 1D, E). However, *Pcdhg^GcKO^* mice made few rears in the open arena and spent proportionally less time exploring the inner zone, which indicate reduced exploration and an anxiety-like phenotype. Similarly in the elevated plus maze, *Pcdhg^GcKO^* mice spent less time and made fewer entries into the open arms, further confirming an anxiety-like phenotype to open space (Figure 1F). *Pcdhg^GcKO^* mutant animals also displayed increased repetitive activity as measured by marble burying (Figure 1G).

*Pcdhg^GcKO^* mutant mice also exhibited spontaneous seizures. To examine seizure activities, cortical electroencephalogram (EEG) recordings were combined with video to monitor wild-type sibling and *Pcdhg^GcKO^* mutant mice (3 months of age, N=4 animals per genotype). We first identified periods of extended immobility and then assessed behavioral seizures using a modified Racine scale (Kim et al., 2019; Racine, 1972). During these periods, *Pcdhg^GcKO^* mutant exhibited seizure symptoms and convulsive behaviors including stiff posturing (scale 1), nodding and forelimb clonus (scale 2, 3) and in a few severe instances, continuous whole body seizures (scale 4) (Figure 1H) (Kim et al., 2019; Racine, 1972). Analysis of simultaneous EEG recordings of baseline cortical activities revealed abnormal bursting and epileptiform discharges during seizure episodes in all mutant *Pcdhg^GcKO^* animals, but not in wild-type (Figure 1I). Together, the neurological and behavioral impairments exhibited by *Pcdhg^GcKO^* mutants suggest significant defects in the development or function of the GABAergic circuitry in the brain.

### GABAergic *Pcdhg* mutants display reductions in brain volume and GABAergic populations

We also observed that juvenile *Pcdhg^GcKO^* mutants developed smaller brains. To locate and quantify the neuroanatomical phenotypes, the brains were dissected at P28 and analyzed by *ex vivo* magnetic resonance imaging (MRI). Compared to control littermates, the total brain volume of *Pcdhg^GcKO^* mice is reduced by 20.5% (Figure 2A,B; N = 12 animals per genotype; *p* <0.0001, Student t-test). Although *Pcdhg^GcKO^* mice are smaller, brain size can vary independently from body size (Allemang-Grand et al., 2017; Ellegood et al., 2013). To identify structure-specific differences, the brains were segmented into 182 regions and measured as percent volume differences from control. Indeed, region-specific volume reductions were detected across the *Pcdhg^GcKO^* brain. The most profoundly affected structures include the globus pallidus, the bed nucleus of stria terminalis, the basal forebrain, and the superior colliculus (Figure 2C). These regions contain high proportions of GABAergic neurons, including interneurons and projection neurons (Kudo et al., 2012; Saunders et al., 2015; Saunders et al., 2016; Villalobos et al., 2018; Whyland et al., 2020). Volume changes were less severe in neocortical areas and the amygdala, which proportionally contain fewer GABAergic neurons (∼20% (Ehrlich et al., 2009; Sahara et al., 2012) (Figure 2C). To verify the correspondence, we performed a correlation analysis of volume reduction detected by MRI and spatial Gad2 expression at E18.5 (as a function of region of interest) obtained from the Allen Brain Institute gene enrichment dataset. The correlation between Gad2 expression and the extent of volume changes in *Pcdhg^GcKO^* indicates that structures with greater developmental Gad2 expression, and thus Cre recombination, are associated with greater volume loss (Figure 2D; −0.267, Fisher-transformed Pearson correlation).

**Figure 2.**
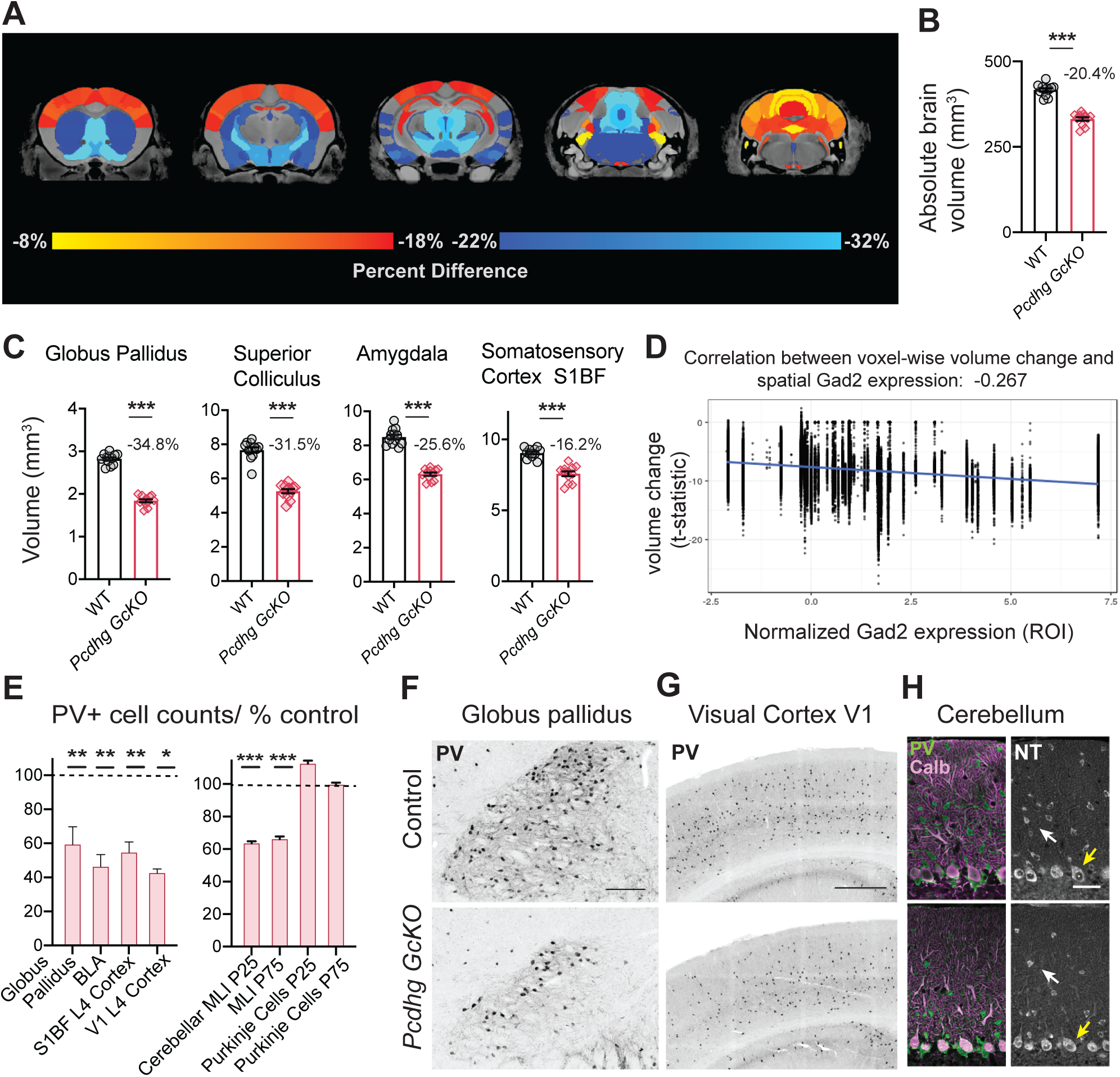
*Pcdhg GcKO* mutants present volume reductions and decreased parvalbumin-expressing populations throughout the brain. **A.** Voxel-wise analysis from whole brain MRI reveals regional reductions in *Pcdhg GcKO* brains at P28, expressed as percent reduction from control brains. On average, the total brain volume significantly smaller than control (−20.4%). Regions coded in blue are further reduced in volume (less than −22%) and those coded in red-yellow are less severely affected (greater than −18%). **B, C.** Absolute volumes of total brain and selected regions of control (black) and *Pcdhg GcKO* (red) brains. Data show mean volume +/− SEM from 12 animals per genotype. All *p* <0.000001, two-tailed Student *t*-test. **D.** MRI volume changes and spatial Gad2 expression at E18.5 across the brain are negatively correlated (−0.267, Fisher-transformed Pearson correlation), indicating association between volume loss severity and spatial Gad2 expression. Voxel-wise volume changes are plotted against spatial Gad2 expression obtained from Allen Brain Institute gene expression dataset. **E.** *Left*, Stereological quantifications of Parvalbumin (PV+) expressing cells in the *Pcdhg GcKO* Globus Pallidus (*p =* 0.0087), Basolateral Amygdala (BLA, *p =*0.0042), Visual cortex V1 layer IV (*p =*0.016), Somatosensory cortex SIBF layer IV (*p =*0.0031). Bars show means from *Pcdhg GcKO* mice normalized as % control sibling data, with % SEM. N=3-4 animals at P28 per genotype, two-tailed Student *t*-test. *Right*, Quantifications of PV+ populations in cerebellar cortex, the Molecular Layer Interneurons (MLI, *p* <0.000001) and Purkinje Cells (PC, *p* = 0.8238) at P25 and P75. N=5-7 animals per genotype. **F, G.** Inverted immunofluorescence images of Parvalbumin-expressing (PV+) cells in Globus Pallidus (F) and Visual cortex (G) in control (top) and *Pcdhg GcKO* (bottom) mice. **H.** *Left*, Immunostaining of PV+ MLIs (green) and PV+/Calbindin+ PCs in cerebellar cortex. *Right*, Soma labeling by Neurotrace. MLI density (white arrow) is reduced but PC density is unchanged (yellow arrow) in *Pcdhg GcKO* cerebella. \**p* < 0.05; ** *p* < 0.01; *** *p* < 0.001. Scale bars: 500µm in F, 200µm in G; 25 µm in H.

The relationship between GABAergic cell populations and volume reduction strongly suggests that the anatomical changes in *Pcdhg^GcKO^* brains result from GABAergic neuron loss. To test this, we began with quantification of neurons immunolabeled for parvalbumin (PV), a Ca^2+^-binding protein that marks subpopulations of GABAergic neurons and interneurons. We first PV-expressing (PV+) neurons in the globus pallidus externus at P28, which comprise ∼55% of the total neuron population and are primarily GABAergic projection neurons that target structures in the basal ganglia, thalamus, and cortex (Hernandez et al., 2015; Kita, 2007). In the globus pallidus of *Pcdhg^GcKO^* animals, the numbers of PV+ neurons are reduced by 40.7% compared to control (Figure 2E, F; N = 4 animals per genotype, *p* = 0.0087, *t-*test). Interneuronal PV+ populations are also diminished, including those in the basolateral amygdala (−53.7%, *p =* 0.0042, *t-*test), and in the primary visual (−53.7%*, p =* 0.0042, *t-*test) and somatosensory barrel cortex (−43.4%, *p* =0.0031, *t-* test) (Figure 2E, G). We confirmed a profound reduction of GABAergic neurons labeled by Gad2-Cre; Rosa26^lsl-tdTomato^ (TdTomato+; −61.5%, *p =* 0.005, *t-*test) in layer IV of the somatosensory cortex, which contains a high density of PV+ interneurons (Sanchez et al., 1992), providing further evidence that *Pcdhg* deletion in GABAergic neurons leads to cell and volume loss (Figure 2 - supplement figure 1). Similarly in the *Pcdhg^GcKO^* cerebellum, the numbers of PV+ molecular layer interneurons were reduced in the cerebellar cortex at P25 (−36.5%, *p* < 0.000001, *t-*test) and did not decline further at P75. PV+ Purkinje cells, which are GABAergic projection neurons, did not differ in *Pcdhg^GcKO^* (Figure 2E, H), consistent with our previous finding that Purkinje-specific *Pcdhg* deletion does not affect Purkinje cell number (Lefebvre et al., 2012). Together, these results demonstrate an essential but cell type-specific role for the Pcdhgs to regulate PV+ GABAergic cell populations throughout the brain, including projection neurons and interneurons.

### Multiple classes of cortical inhibitory interneurons are lost in GABAergic *Pcdhg* mutants

To determine the function of Pcdhgs at the cell-type and developmental levels, we focused on the inhibitory interneuron populations in the somatosensory cortex, which comprise diverse but molecularly well-characterized populations. We first determined whether the Pcdhgs regulate broad or specific classes of GABAergic interneurons by quantifying two cardinal classes of interneurons that originate from the MGE and express PV or Somatostatin (SST+), and two CGE-derived classes that are defined by vasoactive intestinal peptide (VIP+) or by Reelin-positive/SST-negative expression in Layer I (Figure 3A) (Rudy et al., 2011). The densities of CGE-derived Reelin+/SST-and VIP+ interneurons in the *Pcdhg^GcKO^* cortex differed significantly from control (Figure 3B-D). And similarly, the PV+ and SST+ interneuron densities were decreased across all cortical layers (Figure 3E, F). In all cases, there was no accumulation of cINs in particular cortical layers relative to others, indicating that laminar migration is intact in *Pcdhg^GcKO^* mutants. Thus, the Pcdhgs are required for the survival of diverse classes of cortical GABAergic populations.

**Figure 3.**
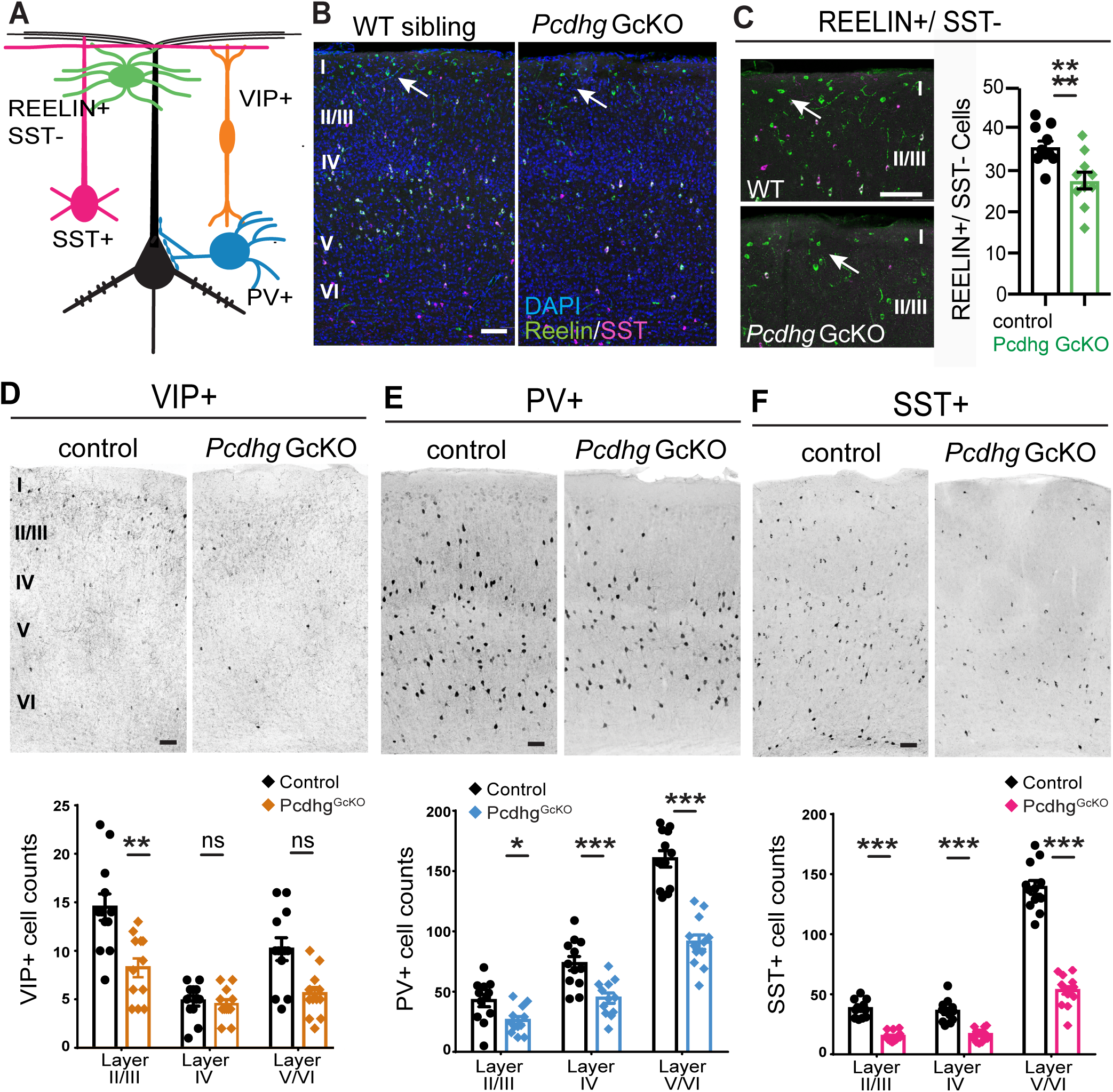
Cardinal classes of cortical GABAergic interneurons are reduced in *Pcdhg* GcKO mutants. **A.** Schematic depicting cortical pyramidal neuron (black) and markers used to distinguish four interneuron (cIN) types. **B-F.** Coronal sections through somatosensory cortex (SSC, SIBF) and quantifications of cIN types at P28 of control and *Pcdhg GcKO* animals, N= 10-12 sections, 3-4 animals per genotype. **B.** Immunostaining for Reelin (green) and SST (magenta) marks Reelin+/SST-population residing in layer I (white arrow). **C.** Labeling and quantification of Reelin+ /SST-cINs in defined area from layer I-III. *p=0.0073* Mann-Whitney (MW) U-test. **D.** Inverted immunofluorescence image (top) and quantifications (bottom) of VIP-expressing cINs. Layers II-III: *p =*0.0015; IV: *p=0.*91; V-VI: *p =* 0.088, MW U-test. **E.** PV-expressing cINs. Layers II-III: *p =* 0.012; IV: *p=*0.00067; V-VI: *p<*0.00001, MW U-test. **F.** SST-expressing cINs. Layers II-III: *p* <0.00001; IV: *p* <0.00001; V-VI: *p<*0.00001, MW U-test. * *p* < 0.05; ** *p* < 0.01; *** *p* < 0.001. Scale bars: 100µm.

### Pcdhgs are required for GABAergic interneuron survival in the cortex and cerebellum during developmentally-regulated cell death

The interneuron loss we observed across hind, mid and forebrain structures of *Pcdhg^GcKO^* brains raises interesting parallels to previous reports of Bax-dependent cell death in early postnatal *Pcdhg* mutant retina and spinal cord (Lefebvre et al., 2008; Prasad et al., 2008). We hypothesized that the Pcdhgs regulate a survival program for GABAergic interneurons in the brain during developmentally-regulated cell death. To test this, we analyzed cortical interneuron PCD which occurs between P5 and P10 and eliminates 30-40% of GABAergic interneurons belonging to the major PV+, SST+, VIP+, 5HTR3/Reelin+SST-classes (Denaxa et al., 2018; Priya et al., 2018; Southwell et al., 2012; Wong et al., 2018). As a first step, we confirmed that *Pcdhgs* are expressed by *Gad2*-positive cINs in postnatal cortex at P7, during the peak of cIN PCD (Figure 4A, B).

**Figure 4.**
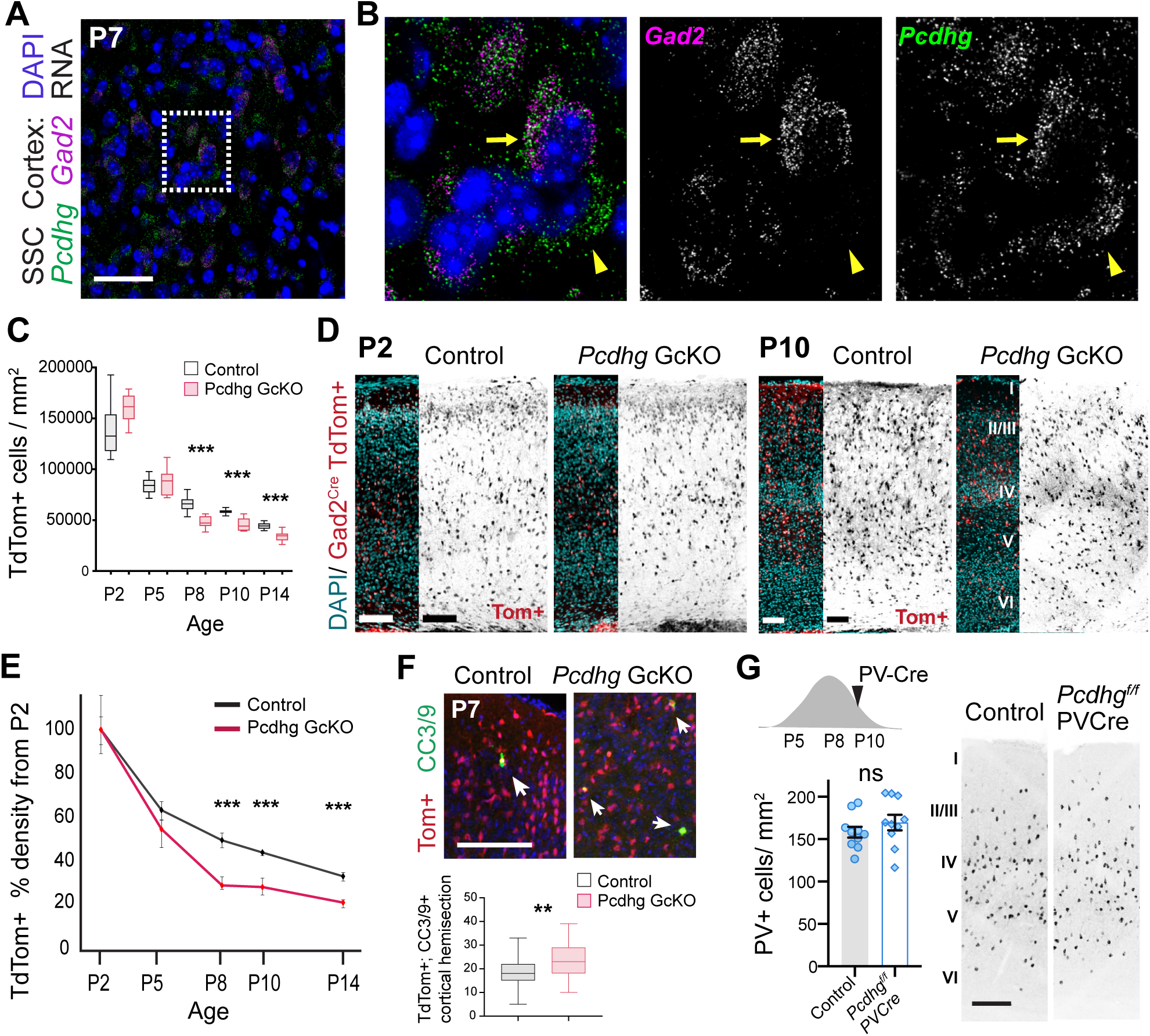
Cortical interneuron loss in *Pcdhg GcKO* mutants is the result of accentuated developmental cell death. **A,B.** Fluorescence *in situ* hybridization of *Gad2* (magenta) and *Pcdhg* (green) mRNAs in P7 SSC cortex. **B.** Inset from A shows *Pcdhg* expression in GABAergic cells (arrow), and in *Gad2-* negative cells (arrowhead). **C.** Quantification of cIN density through development in control (grey) and *Pcdhg GcKO* SSC cortex (red). cINs are marked by Gad2-Cre; Ai14-TdTomato. N=12 sections, 3 animals per genotype. P8, *p =* 0.000005; P10, *p =* 0.0003; P14, *p =* 0.000009, Mann-Whitney U-test. **D.** Coronal section of control and *Pcdhg GcKO* SSC at P2 and P10 with labeling of nuclei (DAPI, cyan) and TdTomato+ cINs (red). Inverted fluorescent images of TdTomato+ cINs are on right. **E.** Graphs show data of TdTomato+ cIN densities from C, normalized to densities at P2 for control (black) and *Pcdhg GcKO* (red). **F.** *Top*, apoptotic TdTomato+ cINs (red) co-labeled with cleaved caspase-3 and −9 (green; white arrows). *Bottom*, quantifications of TdTomato/cleaved caspase-3/9 cells in P7 cortical sections. N=31 sections, from 3 animals per genotype; *p=* 0.00095, Mann-Whitney U-test. **G.** *Top left*, the onset of PV-Cre activity occurs after the peak of cIN PCD. *Bottom left*, PV+ densities of SSC at P28 are indistinguishable between control (grey bars) and *Pcdhg ^f/f^; PV-Cre* animals (white). Bars show mean +/− SEM from 10 sections, 3 animals per genotype. *p* = 0.32, unpaired *t-*test. *Right*, coronal sections with PV+ labeling of SSC of control and *Pcdhg ^f/f^; PV-Cre* animals. * *p* < 0.05; ** *p* < 0.01; *** *p* < 0.001. Scale bars: 50µm in A; 100µm in D-F; 200µm in G.

To determine whether cIN loss in *Pcdhg^GcKO^* mutants results from dysregulated PCD, the numbers of cINs were quantified in control and mutant *Pcdhg^GcKO^*; Rosa^LSL-TdTomato^ cortices during postnatal development. From P2 through P14, the densities of TdTomato+ cINs declined in both control and *Pcdhg^GcKO^* mutants, consistent with the postnatal timecourse of cIN loss to PCD (Southwell et al., 2012; Wong et al., 2018) (Figure 4C-4E). Early in postnatal development, TdTomato+ cIN densities in mutant *Pcdhg^GcKO^* cortex were indistinguishable from those in control (Figure 4C-D), indicating that similar numbers of cINs are born and migrate to the cortex. By P8, the TdTomato+ cIN densities declined to a greater extent in *Pcdhg^GcKO^* mutant and were reduced by 25% compared to controls by P14 (Figure 4E). To compare the extent of cell death in *Pcdhg^GcKO^* mutants from naturally-occurring apoptosis in controls, we quantified apoptotic TdTomato+ cINs marked by cleaved caspase-3 and caspase-9 at P7 and confirmed a complementary increase in apoptosis in *Pcdhg^GcKO^* cortex (Figure 4F). Thus, the excessive loss of cINs in *Pcdhg^GcKO^* mutants occurs due to increased postnatal apoptosis. To test if the survival requirement for the Pcdhgs is temporally restricted to this postnatal period, *Pcdhgs* were deleted from cINs after the peak of PCD using Parvalbumin-ires-Cre, which begins to be expressed in maturing PV+ cINs after the first postnatal week (Lazarus et al., 2015). In this experiment in which the *Pcdhg* genes remain largely during peak PCD, the PV+ cell densities in the cortex of *Pcdhg^f/f^; PVCre* animals are indistinguishable from control (Figure 4G).

In a final test to show that regulation of cIN number by the Pcdhgs is specific to PCD rather than other degenerative processes, we tested whether cIN loss in *Pcdhg^GcKO^* mutants occurs through the *Bax-*dependent apoptotic pathway. In *Bax* mutant mice, genetic blockade of apoptosis prevents the natural elimination of cINs (Priya et al., 2018; Southwell et al., 2012; Wong et al., 2018). If excess cIN loss in *Pcdhg^GcKO^* cortex results from the overactivation of apoptosis, then the cINs destined to die in *Pcdhg^GcKO^*; *Bax* double mutants (*Pcdhg^f/f^; Bax ^f/f^; Gad2Cre*) would be preserved resulting in greater numbers of mature cINs. Indeed, the PV+ and Reelin+ cIN populations in *Pcdhg^GcKO^; Bax^GcKO^* double mutant animals were not only restored compared to *Pcdhg^GcKO^* single mutants but were also indistinguishable from control *Pcdhg^Het^; Bax^GcKO^* siblings (Figure 5A, B). Together, this series of experiments establish an essential role for the Pcdhgs to promote cIN survival during the period of PCD.

**Figure 5.**
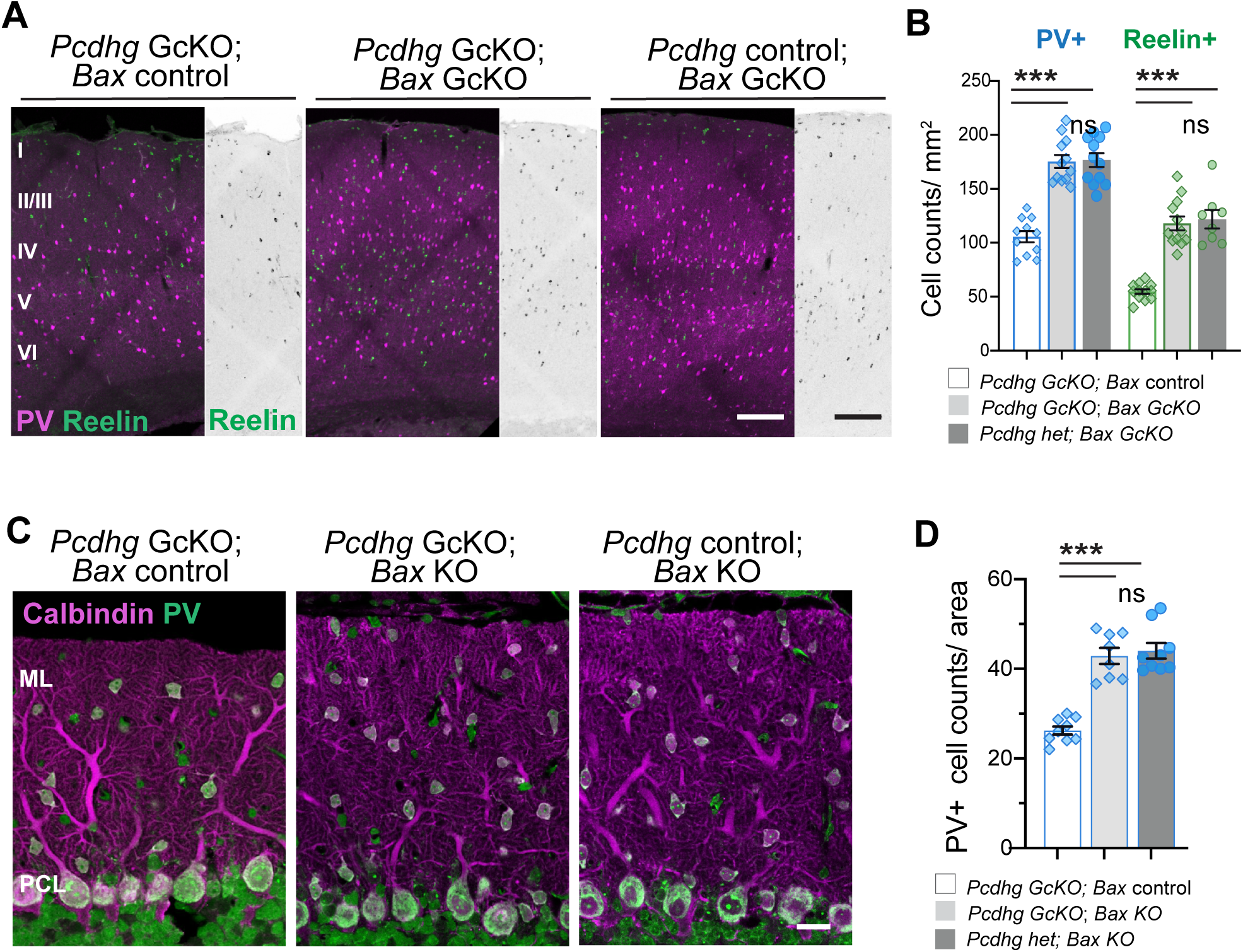
Genetic blockade of apoptosis by *Bax* deletion rescues cortical and cerebellar interneuron loss in *Pcdhg GcKO* mutants. **A.** Immunostaining for PV (magenta) and Reelin (green) in SSC of *Pcdhg GcKO, Pcdhg GcKO; Bax GcKO*, and *Bax GcKO* animals. Inverted images of Reelin+ labeling are shown on right. **B.** Quantifications of PV+ and Reelin+ cINs across cortical layers show rescue in *Pcdhg GcKO; Bax GcKO* double mutants compared to *Bax GcKO* control animals (ns, not significant, *p >* 0.99). Bars show mean +/− SEM and counts from 12 sections, 4 animals per genotype. PV+: *Pcdhg GcKO* compared to *Pcdhg GcKO; Bax GcKO*, *p =* 0.00015; and to *Pcdhg* control*; Bax GcKO, p =* 0.00011. Reelin+: *Pcdhg GcKO* compared to *Pcdhg GcKO; Bax GcKO*, *p =* 0.00012; and to *Pcdhg* control*; Bax GcKO, p =* 0.00039. **C.** Immunostaining of molecular layer interneurons (MLI) labeled by PV (magenta) and Neurotrace (NT, green) in cerebellar cortex of *Pcdhg GcKO, Pcdhg GcKO; BaxKO*, and *BaxKO* mutant animals. **D.** Quantifications show rescue of cerebellar PV/NT+ interneurons in *Pcdhg GcKO; BaxKO* double mutants compared to *BaxKO* control animals (ns, not significant, *p >* 0.99). Bars show mean +/− SEM and counts from 8-9 sections, 3 animals per genotype. *Pcdhg GcKO* compared to *Pcdhg GcKO; BaxKO*, *p =* 0.00031; and to *Pcdhg* control*; BaxKO, p =* 0.000057. Kruskal-Wallis multiple comparison tests. *** *p* < 0.001. Scale bars: 200µm in A; 20µm in C.

We next sought to address whether the pro-survival role for the Pcdhgs in PCD extends to GABAergic interneuron populations elsewhere in the brain and that differ in embryonic origin. We focused on the PV+ cerebellar molecular layer interneurons that originate from a *Ptf1a*-expressing GABAergic lineage in the ventricular zone of the cerebellar primordium (Hoshino et al., 2005; Leto et al., 2006). Between P5 and P12, these interneurons normally undergo apoptosis in the cerebellar prospective white matter, where precursors proliferate and differentiate, and in the cerebellar cortex (Sergaki et al., 2017; Yamanaka et al., 2004). Apoptosis of the molecular layer interneurons is also Bax-dependent (Figure 5 – figure supplement 1A). Similar to forebrain cINs in *Pcdhg^GcKO^* animals, increased caspase-3 positive signals were detected in the *Pcdhg^GcKO^* cerebellar cortex at P7 (Figure 5 – figure supplement 1B). Moreover, in double *Pcdhg^GcKO^*; *Bax^KO^* mutant cerebellum, the density of PV+ positive interneurons is restored and indistinguishable from *Bax^KO^; Pcdhg* control siblings (Figure 5C, D). By contrast, *Pcdhg*- and *Bax*-dependent regulation of PCD does not require p53, a transcriptional activator of several pro-apoptotic genes, because conditional deletion of *Tp53* gene with Gad2-Cre does not cause ectopic survival of the molecular layer interneurons (Figure 5 – figure supplement 2).

Collectively, our results demonstrate that the Pcdhgs promote the survival of two unrelated GABAergic interneuron populations during postnatal PCD. We propose that the Pcdhgs act broadly in multiple GABAergic interneuron populations in the brain to regulate neuronal survival during PCD.

### Pro-survival Akt-FoxO3A signals are diminished in *Pcdhg* mutant GABAergic interneurons during PCD

Whether developing neurons survive or die is determined by the balance of intracellular signals that suppress or initiate the apoptotic cascade (Hollville et al., 2019; Pfisterer and Khodosevich, 2017). To assess how the Pcdhgs influence this balance, we evaluated the activity of intracellular effectors that suppress intrinsic apoptosis, including the serine-threonine kinase AKT which is elevated in activity during cIN PCD (Wong et al., 2018). Differences in phospho-AKT levels between control and *Pcdhg^GcKO^* cortical lysates at P7 could not be distinguished by standard Western blots (Figure 6 – supplement figure 1A). To discriminate cIN-specific changes, Western blots were performed on dissociated TdTomato-labeled cINs isolated by FACS. In cIN-enriched P7 cell lysates, phospho-AKT levels were detectably lower in *Pcdhg^GcKO^* samples (Figure 6 – Supplement Figure 1B) but quantifications were hampered by the low amounts of material obtained. We therefore applied a quantitative approach using flow cytometry to measure phospho-AKT levels in large numbers of dissociated cINs labeled by Gad-Cre;TdTomato-labeled cINs. As a population, the *Pcdhg^GcKO^* cINs exhibited significant reductions in phospho-AKT signals compared to control cINs. Therefore, AKT signaling is diminished in developing *Pcdhg*-deficient cINs (Figure 6A) during the period of PCD.

**Figure 6.**
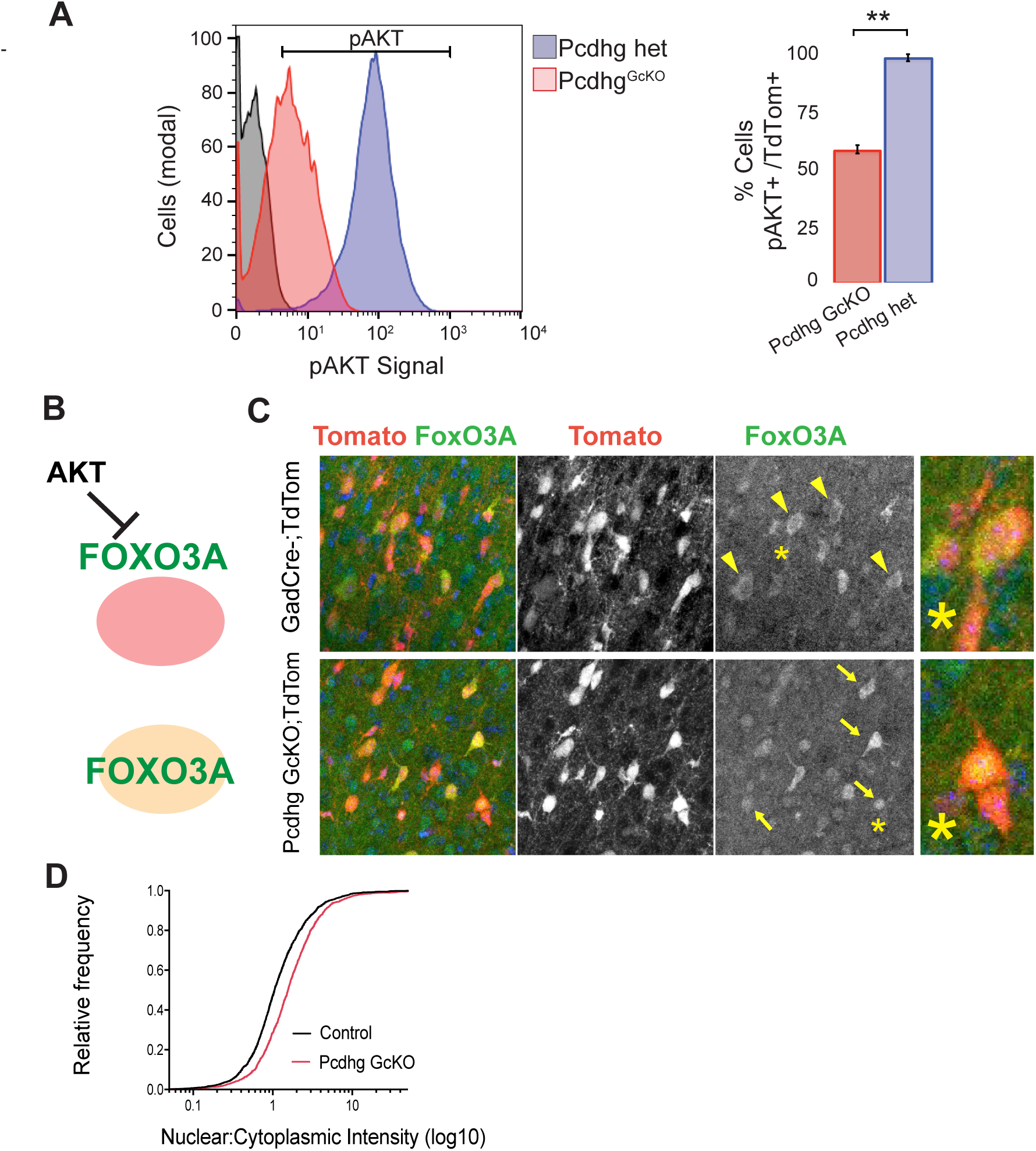
Pro-survival Akt-Foxo3A signaling is diminished in Pcdhg GcKO mutants. **A.** *Right*, Cytometric measure of pAKT immunolabeled Tdtomato+ cortical cells isolated from *Pcdhg GcKO* (GcKO; red) and heterozygous Pcdhg control (het; blue) p7 cortical dissociations. Left panel depicts decreased pAKT immunofluorescence in the Pcdhg GcKO cell population relative to Pcdhg controls. Antibody specificity control (black). *Left*, Histogram of pAKT immunopositive cells plotted as percentage of Tdtomato+ cells population (N = 4 animals per genotype; p = 0.0032; two-tailed t-test). **B.** Schematic of FoxO3A translocation from cytoplasmic (top) to nuclear (bottom) compartments in the absence of AKT-dependent inhibition. **C.** Immunostaining of P8 cortex shows translocation of FoxO3A (green) in Gad2Cre-TdTomato+ cINs (red) from a predominantly cytoplasmic localization in control (top, yellow arrowheads) to nuclear enrichment in *Pcdhg GcKO* mutants (bottom, yellow arrows). **D.** High magnification of inset (*) from **C**. **D.** FoxO3a fluorescence intensity in the nucleus and cytosol of control (black) and *Pcdhg GcKO* (red) TdTomato+ cINs across layers II-VI were quantified and plotted as a nuclear to cytoplasmic ratio. Graph shows cumulative distribution and increased nuclear to cytoplasmic ratio in *Pcdhg GcKO*. N = 3 animals per genotype, *p* <0.00001, Kolmogorov-Smirnov test. *** *p* < 0.001.

One mechanism by which AKT is anti-apoptotic is through phosphorylation and cytoplasmic retention of the FoxO transcription factors and thus prevention of FoxO-mediated transcription of targets such as pro-apoptotic BH3 members *puma* and *bim* (Figure 6B) (Ambacher et al., 2012; Gilley et al., 2003). We predicted that diminished AKT signaling in *Pcdhg^GcKO^* cINs would result in increased FoxO3A translocation to the nucleus and upregulation of *puma* and *bim* mRNA. We compared FoxO3A activity in *Pcdhg^GcKO^* and control cINs by quantifying the ratios of nuclear (active) and cytoplasmic (inactive) FoxO3A protein accumulation by immunofluorescence. During the period of PCD, FoxO3A is noticeably enriched in the cytoplasm of TdTomato-labeled cINs in control. In the *Pcdhg^GcKO^* cortex, FoxO3A signal is higher in the nucleus, as confirmed by the increased nuclear-to-cytoplasmic ratio of FoxO3A+ fluorescence (Figure 6C, D). Based on these results, we conclude that the Pcdhgs promote cIN survival through a mechanism involving modulation of the AKT pathway.

### cIN survival does not require Pcdhg-dependent interactions from other pyramidal or cIN neurons

As a large family of transmembrane receptors with homophilic binding properties, the Pcdhgs are interesting candidates to mediate cell-cell interactions for selective cIN survival. To begin to dissect these relationships, we used genetic approaches to test whether cIN survival requires coordinated Pcdhg expression in other cortical cell populations: 1) pyramidal neurons, the major synaptic partners that collectively serve as cIN afferents and targets; 2) cortical astrocytes; 3) MGE-derived cINs to determine if they influence the size of CGE populations; and 4) mosaically distributed Pcdhg^WT^ and Pcdhg^GcKO^ cINs belonging to same type.

To test whether Pcdhgs in principal neurons contribute to cIN survival, *Pcdhg^f^* mice were crossed to Emx1-Cre to delete *Pcdhgs* from forebrain progenitors that give rise to pyramidal cells and glia, but spare the *Pcdhg* genes in the ganglionic-eminence derived cINs (Figure 7A). The survival of pyramidal neurons is not affected in *Pcdhg^f/f^*; Emx1-Cre mutant mice (Garrett et al., 2012). The densities of mature PV+, SST+ and Reelin+/SST-cIN populations were also unaffected in *Pcdhg^f/f^*; Emx1-Cre mutants, demonstrating that appropriate levels of PCD occur in the absence of *Pcdhgs* in pyramidal neurons (Figure 7B-D). Therefore, cIN survival does not require Pcdhg interactions from the pyramidal populations nor cortical glia.

**Figure 7.**
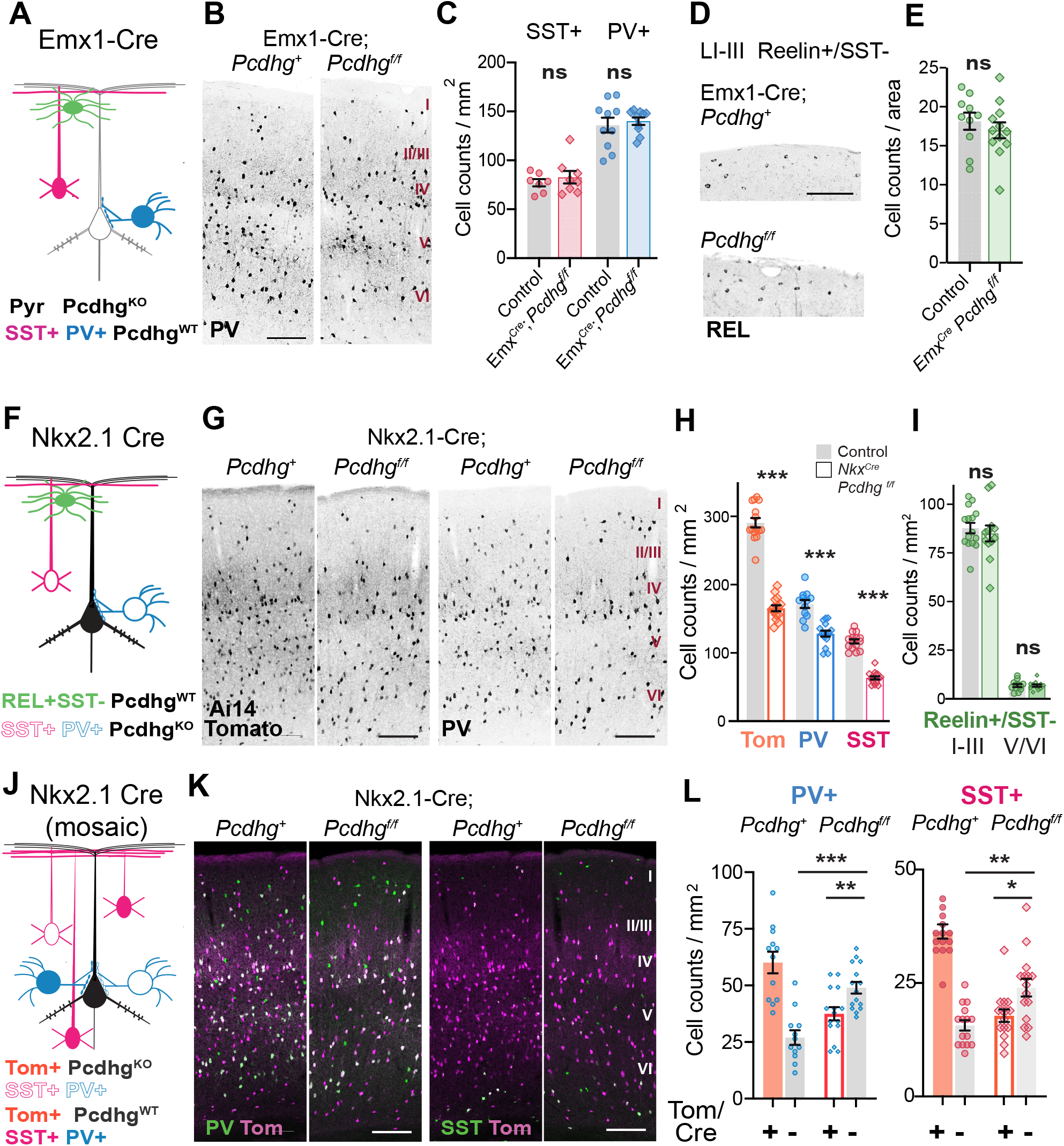
Cell autonomous requirement for Pcdhgs for cIN survival. **A.** Test for Pcdhgs in Pyramidal populations: Emx1-Cre deletes *Pcdhgs* from pyramidal and glial cells, leaving the Pcdhgs intact in cINs. **B.** Inverted fluorescent image of PV+ cINs in control and *Pcdhg ^f/f^;Emx1-Cre* mutants at P28. **C.** Quantifications show that PV+ (*p* = 0.65, two-tailed *t-test*) and SST+ cINs (*p=*0.48) are significantly different (ns) in *Pcdhg ^f/f^;Emx1-Cre* mutants compared to control. **D.** Inverted image of Layer I CGE-Reelin+ cINs in control and *Pcdhg ^f/f^;Emx1-Cre* mutants. **E.** Quantifications of Reelin+/SST-cINs in layer I-III in control and *Pcdhg ^f/f^;Emx1-Cre* mutants (shaded green). Data in C, E show mean +/− SEM from 12-15 sections, 3-4 animals per genotype, *p* = 0.6, two-tailed *t-test*. **F.** Test for non-autonomous role in CGE-derived Reelin+/SST-(green), using Nkx2.1-Cre to delete *Pcdhg^f/f^* from MGE-derived PV+ (blue) and SST+ (pink) cINs. **G.** Inverted fluorescent images of Ai14Tomato+ (left) and PV+ (right) cINs of control and *Pcdhg^f/f^;Nkx2.1Cre;Ai14-LSL-TdTomato* animals. **H.** Quantifications of MGE-derived Tomato+ (*p* <0.000001, two-tailed *t-*test), PV + (*p*=0.000002, two-tailed *t-*test), and SST+ (*p* <0.000001, two-tailed *t-*test) populations in control (grey bar) and *Pcdhg ^f/f^;Nkx2.1Cre* mutants (outlined bar). **I.** Quantifications of Reelin+/SST-cINs (green) in layers I-III (*p* = 0.71) and layers V-VI (*p* =0.99) by two-tailed *t-test*). Data in H, I show mean +/− SEM from 12-15 sections, 3-4 animals per genotype. **J.** Test for intrinsic role by comparing Pcdhg^WT^ (Nkx2.1Cre-; Tomato-) and Pcdhg^KO^ (Nkx2.1Cre+; Tomato+) PV+ and SST+ cINs using Nkx2.1-Cre which has mosaic activity. **K.** Immunofluorescent images of Tomato (magenta) with PV+ (right, green), or Tomato (magenta) with SST+ (left, green) populations in control and *Pcdhg ^f/f^;Nkx2.1Cre;Ai14Tomato* animals. Note prevalence of Green+/Tomato-cells. **L.** Quantifications of PV and SST cINs in Layers I-III marked by Cre+/Tomato+ or Cre-/Tomato-subpopulations. In *Pcdhg ^f/f^;Nkx2.1Cre;Ai14Tomato* animals, there are significantly more Cre-/Tomato-cINs (grey bar) compared to Cre+/Tomato+ (orange outline) (PV+: *p =*0.0064; SST+: *p=*0.013, Mann-Whitney test) and to control Cre-/Tomato-cINs (PV+: *p =*0.00001; SST+: *p=*0.001, Mann-Whitney test). Data show mean +/−SEM from 15 sections from 3-4 animals per genotype. * *p* < 0.05; ** *p* < 0.01; *** *p* < 0.001. Scale bars: 200µm.

A compensatory relationship has been proposed between MGE- and CGE-derived cINs, in which MGE-cIN reductions lead to increased survival of CGE-derived 5HTR3+/Reelin+ interneurons, particularly in the deeper layers where this population is normally scarce (Denaxa et al., 2018). To test whether the Pcdhgs influence the survival of CGE-cIN non-autonomously, we used Nkx2.1-Cre to delete *Pcdhgs* from the MGE lineage (Figure 7F). As expected, the total Nkx-Cre; tdTomato+ cINs as well as the SST+ and PV+ populations are significantly reduced in *Pcdhg^f/f^*; Nkx2.1-Cre mutants (−43.1%, −45.7%, and −25.5%, respectively) (Figure 7G-H). However, the numbers of CGE-derived Reelin+/SST-cINs are indistinguishable from those in control (Figure 7I) in the upper and lower layers. Thus the size of the Pcdhg-expressing CGE-cIN population is not influenced by *Pcdhg-*dependent loss of MGE-cINs (Figure 7I).

Based on our observation that *Pcdhg* mutant cINs have diminished pro-survival signals, we tested whether wild-type cINs (Pcdhg^WT^) have a competitive advantage over sibling Pcdhg^KO^ cINs to survive and incorporate into the developing cIN network. As confirmed by Cre-dependent TdTomato labeling (Figure 7 - supplement figure 1), we took advantage of the mosaic recombination activity of Nkx2.1-Cre (Xu et al., 2008) in the upper cortical layers to test if the proportions of surviving Pcdhg^WT^ PV+ and SST+ cINs are positively or negatively changed in *Pcdhg^f/f^*; Nkx2.1-Cre mutant layers (Figure 7J). Interestingly, the numbers of Pcdhg^WT^ PV+; Tomato-negative and Pcdhg^WT^ SST+; Tomato-negative cINs are significantly increased in LI-III in *Pcdhg^f/f^*; Nkx2.1-Cre mutants at P28 (Figure 7K,L; Figure 7 – supplement figure 1), suggesting that proportionally more Pcdhg^WT^ cINs survived the period of PCD compared to *Pcdhg^f/f^*; Nkx2.1-Cre positive cells. Based on the selective survival of Pcdhg^WT^ cINs over Pcdhg^KO^ cells, we conclude that the Pcdhgs mediate an intrinsic advantage in cINs to survive and establish in the maturing cortex.

## DISCUSSION

The sculpting of inhibitory interneuron populations is critical for the formation and function of neural circuits. In the neocortex, the elimination of GABAergic interneurons by PCD must be paired with mechanisms that select optimal numbers and qualities of interneurons for survival and integration. Similar programs likely operate throughout the developing brain to sculpt the GABAergic circuitry. In this study, we determined that the family of gamma-Protocadherin (Pcdhgs) receptors regulates a pro-survival mechanism in cortical GABAergic interneurons during postnatal PCD. In GABAergic *Pcdhg* mutants, cINs are initially established at similar densities in the neonatal cortex but decline excessively due to accentuated Bax-dependent apoptosis. By deleting the Pcdhgs in cortical subpopulations, we determined that cIN survival does not depend on the Pcdhgs in pyramidal cells but results from an enhanced pro-survival capacity due to intrinsic Pcdhg expression. The Pcdhgs regulate a similar mechanism of interneuron survival in the cerebellum and they are essential for the development and function of the GABAergic circuitry throughout the brain.

### The Pcdhgs mediate a pro-survival mechanism for GABAergic interneuron PCD

Developmental cell death of GABAergic interneurons in the cortex is a process that has been characterized recently (Denaxa et al., 2018; Priya et al., 2018; Southwell et al., 2012; Wong et al., 2018). These studies established that cIN PCD occurs through the intrinsic, Bax-dependent apoptotic pathway, and is influenced by activity-dependent and –independent factors. We propose that the Pcdhgs serve as a molecular mechanism for GABAergic interneuron survival during the process of PCD, and discuss how key findings here build on previous reports.

First, the Pcdhgs are required for the survival of cINs belonging to the four major MGE- and CGE-derived classes shown to undergo PCD, and therefore not selective to subpopulations influenced by activity-dependent or independent mechanisms (Denaxa et al., 2018; Priya et al., 2018; Southwell et al., 2012). The excitatory pyramidal cells do not depend on the Pcdhgs for their survival (Garrett et al., 2012). Similarly in the cerebellum, the Pcdhgs have a pro-survival role for GABAergic interneurons but not for the Purkinje cells, which are GABAergic projection neurons. Cell type-selectivity is a key feature of Pcdhg-dependent survival but principal neurons are not entirely excluded. For instance, the Pcdhgs regulate the survival of principal neurons in the retina, the retinal ganglion cells (Ing-Esteves et al., 2018; Lefebvre et al., 2008), but are dispensable for motorneurons in the spinal cord (Hasegawa et al., 2016; Prasad et al., 2008; Wang et al., 2002). In the brain, Pcdhg-dependent survival might extend to subsets of GABAergic projection neurons that undergo PCD such as the globus pallidus (Waters et al., 1994) which is severely reduced in size in *Pcdhg^GcKO^* mutants, and in the hypothalamus (Kong et al., 2012; Su et al., 2010).

Second, the regulation of cIN survival by the Pcdhgs follows the natural timeline of cIN PCD (Denaxa et al., 2018; Priya et al., 2018; Southwell et al., 2012; Wong et al., 2018). As demonstrated in heterochronic transplantations, the timing of cIN PCD is intrinsically-determined. Grafted cINs undergo apoptosis according to their developmental stage rather than the timing of PCD of the host circuitry (Southwell et al., 2012). Our results also indicate that the responsiveness of cINs to pro-survival Pcdhgs is restricted to this period because Pcdhg deletion after PCD does not cause further death. Thus, the Pcdhgs regulate the extent but not the timing of cIN PCD. The accentuated death of spinal interneurons in *Pcdhg* mutants also follows the exquisite, cell type-specific ordering of PCD (Prasad et al., 2008). The MGE-derived cIN wave of PCD slightly precedes that of the CGE-cINs (Priya et al., 2018), suggesting that cIN susceptibility to cell death is tuned to birth date and cell type. Together, these studies indicate that cINs are temporally-restricted to respond to pro-survival or pro-death signals and to regulate the apoptotic cell death machinery accordingly. Soon after, pro-apoptotic factors are likely downregulated to terminate the wave of PCD (Hollville et al., 2019) and to stabilize final cIN populations for circuit maturation. Interestingly in the visual cortex, subpopulations of MGE-derived chandelier cells succumb to a second wave of apoptosis for additional pruning to match binocular inputs and enable experience-dependent refinement necessary for binocular vision (Wang et al., 2019). Thus, developmental PCD might be implemented for multiple tasks to assemble and refine specialized circuits.

Finally, we propose that the Pcdhgs promote cIN survival through modulation of AKT signaling, which is elevated during the peak of cIN PCD (Wong et al., 2018). Identifying signals downstream of the Pcdhgs that positively regulate survival has been challenging due to the cell type-specificity and lack of *in vitro* model systems that recapitulate Pcdhg-dependent survival. By quantifying intracellular components in purified cIN populations during PCD, we detected a reduction in phospo-AKT levels in *Pcdhg^GcKO^* mutant cINs. The PI3K-AKT pathway is a prominent regulator of neuronal survival through multiple anti-apoptotic and pro-survival actions (Brunet et al., 2001). One link between diminished AKT signaling and increased apoptosis in *Pcdhg^GcKO^* cINs is the translocation of FOXO3A, which upregulates pro-apoptotic factors such as *Puma* and *Bim* (Ambacher et al., 2012; Li et al., 2009). The relationship between AKT and Pcdhgs is especially interesting because PI3K-AKT regulates cortical and hippocampal neuron survival by integrating numerous upstream signals, including trophic factors, spontaneous depolarization and synaptic transmission, and intracellular calcium signaling (Golbs et al., 2011; Hardingham et al., 2002; Heck et al., 2008; Leveille et al., 2010; Murase et al., 2011; Wagner-Golbs and Luhmann, 2012). Our finding that wild-type cINs are more likely to survive than *Pcdhg* mutant cINs in a mosaic *Pcdhg* cIN population (Figure 8) also underscores the intrinsic importance of Pcdhgs. One potential mechanism is that variations in Pcdhg-AKT signaling among cINs contribute to the threshold for survival or cell death, and provide differential advantage resulting in selective cell survival.

Further studies are required to identify signals that are recruited by the Pcdhgs at the cell membrane. One pro-apoptotic candidate is PTEN, which negatively regulates PI3K and is required for cIN PCD (Wong et al., 2018). PYK2 and FAK are negatively regulated by the cPcdhs for pyramidal neuron migration and arborization (Fan et al., 2018; Garrett et al., 2012), and overexpression of PYK2 induces apoptosis in chick spinal cord (Chen et al., 2009). Studies are underway to test these candidates. The intracellular Pcdhg domains required for survival are unknown. The 22 Pcdhg isoforms share an identical cytoplasmic terminus but bear unique juxtamembrane domains; both domains regulate Pcdhg trafficking (Fernandez-Monreal et al., 2009; O’Leary et al., 2011; Shonubi et al., 2015). Pcdhg-dependent survival will involve isoform-specific signalling through extracellular and juxtamembrane regions because the subset of C-type Pcdhg genes, and PcdhgC4 in particular, is essential for neuronal survival (Chen et al., 2012; Garrett et al. 2019). Diverse cPcdhs isoforms might also signal together as Pcdh-alpha and Pcdh-beta members multimerize and functionally cooperate with the Pcdhgs to contribute to neuronal survival (Goodman et al., 2017; Goodman et al., 2016; Han et al., 2009; Hasegawa et al., 2016; Ing-Esteves et al., 2018). Also, developmental survival of cerebellar interneurons is regulated by GDNF receptors RET and GFR1a (Sergaki et al., 2017), raising the interesting possibility of co-signaling through interactions between RET and Pcdh-alpha members (Schalm et al., 2010).

### Pcdhg-dependent interactions and cIN survival

The purpose of cIN developmental death is presumed to numerically or functionally match inhibitory interneurons to the excitatory pyramidal populations (Wong and Marin, 2019). PCD might also adjust the size of inhibitory neural assemblies through local, population-intrinsic mechanisms (Southwell et al., 2012) or network mechanisms (Duan et al., 2019). As a large family of transmembrane receptors with homophilic binding properties, the cPcdhs are enticing candidates for differential cIN survival through selective cell-cell interactions. Given the potent effects of manipulations of pyramidal cell activity or number on cIN survival (Wong and Marin, 2019), one attractive hypothesis is that the cPcdhs underlie cell-cell or synaptic interactions between the cINs and pyramidal neurons. We tested the simplest scenario: whether interneuronal survival requires Pcdhg expression in the principal neurons providing afferent and target connectivity. But we found that Pcdhgs are not required in pyramidal neurons, ruling out a target/afferent-dependent requirement for Pcdhgs in cIN survival. This aligns with findings from retina where synaptic targeting and formation are not prerequisites for Pcdhg-dependent survival (Lefebvre et al., 2008), though the Pcdhgs influence both synaptic contacts and survival in the spinal cord (Chen et al., 2012; Prasad et al., 2008; Weiner et al., 2005). However, our results do not account for the potential redundancy among other cPcdh members, or other ways in which cPcdh complexes might interact with RTKs, neurotransmitter receptors, or synapse-organizing molecules (Li et al., 2012; Molumby et al., 2017; Schalm et al., 2010). Although our experiments do not distinguish between activity-dependent or independent actions, the Pcdhgs are required for VIP+ cIN survival, which is not influenced by activity (Denaxa et al., 2018; Duan et al., 2019; Priya et al., 2018).

A second possibility is that the Pcdhgs mediate interactions among the cINs to enhance electrical coupling and synchronized network activities, which emerge during the first postnatal week (Duan et al., 2019; Minlebaev et al., 2011; Pangratz-Fuehrer and Hestrin, 2011). cPcdh deletions result in abnormal network activities in spinal cord and synchronized activities *in vitro* (Hasegawa et al., 2017), suggesting possible roles in patterning network topology. As demonstrated recently (Duan et al., 2019), network activities that influence cIN survival are complex and dependent on GABAergic signaling onto pyramidal neurons (Duan et al., 2019). Understanding how the complex group of Pcdhgs sculpts interneuronal populations requires mechanistic investigations of Pcdhg functions at the molecular, cellular and circuit level.

### The Pcdhgs are required for proper development of the GABAergic circuitry

In addition to detailed characterizations of Pcdhgs in the cortical and cerebellar interneurons, our results highlight the broad importance for the Pcdhgs in the development and size of GABAergic neuron populations throughout the brain. In brains of juvenile GABAergic *Pcdhg* mice, structures most notably reduced in volume include regions with proportionally higher GABAergic content, such as in basal ganglia and the midbrain. The behavioral deficits and seizures exhibited by GABAergic *Pcdhg* mutant mice point to broad GABAergic defects and further demonstrate the importance of cPcdhs in complex CNS patterning and behaviors (Chen et al., 2017; Fukuda et al., 2008; Yamagishi et al., 2018). Cell type- and developmental-specific deletions are required to dissect additional roles for the Pcdhgs in GABAergic neuronal patterning, branching and maturation after PCD.

In conclusion, we identify the Pcdhgs as an essential regulator of GABAergic interneuron survival in the developing brain. We propose a model in which the extent of PCD among cINs is determined by pro-survival Pcdhg-AKT signaling to provide cINs the advantage to survive and integrate. As variants and dysregulated expression of the human PCDH gene cluster are associated with neurodevelopmental and psychiatric disorders including ASD, Tourette’s, and schizophrenia (Anitha et al., 2013; El Hajj et al., 2016; Iossifov et al., 2012; Schizophrenia Working Group of the Psychiatric Genomics, 2014; Shao et al., 2019; Vadodaria et al., 2019), these findings also have relevance for understanding the developmental basis of inhibitory deficits in human disorders.

## METHODS

### Mouse Strains

Mouse lines used in this study have been described previously. The *Pcdhg ^fcon3^* conditional allele contains loxP sequences that flank the third constant exon and generates a functionally null allele following Cre recombination (Lefebvre et al., 2008; Prasad et al., 2008). The *Gad2-ires-cre* knock-in line (JAX #028867) co-expresses *Cre* with *Gad2/*Gad65 in the majority of developing GABAergic neurons and a small percentage of cortical astrocytes (Taniguchi, 2014; Taniguchi et al., 2011). The constitutive and conditional floxed *Bax* alleles were used for the cerebellar and neocortical studies, respectively (*Bax^tm1Sjk^*, JAX # 002994; *Bax^tm2Sjk^*, JAX# 006329) (Knudson et al., 1995; Takeuchi et al., 2005). In contrast to the available *Bax^tm2Sjk^ Bak1^tm1Thsn^* mice, the knockout *Bak* allele was not maintained with *Bax ^f/f^* for this study. The following lines have also been described: *Emx1-ires-cre* (JAX #005628) (Gorski et al., 2002); *Nkx2.1-cre* BAC transgenic (JAX #008661) (Xu et al., 2008); *PV-ires-cre* (Pvalbtm1(cre)^Arbr/J^; JAX #017320) (Hippenmeyer et al., 2005); the tdTomato Cre reporter line Ai14-RCL-tdT (JAX# 007914) (Madisen et al., 2010); *Tp53* floxed mice (JAX# 007914) (Marino et al., 2000). Mice were maintained on a C57/B6J background. For all analyses in this study, wild-type and heterozygous *Pcdhg;* Cre positive siblings were selected for the control cohorts, and mice of both sexes were used. Animals were coded to blind the investigators of the genotypes.

All animal experiments were performed in accordance with Canadian Council on Animal Care Guidelines for Use of Animals in Research and Laboratory Animal Care under protocols approved by The Centre for Phenogenomics Animal Care Committee and by the Laboratory of Animal Services Animal Use and Care Committee at the Hospital for Sick Children (Toronto, Canada).

### Behavioral Assays

#### Open-Field Test

Locomotor activities of the mice were examined in a clear square Plexiglass apparatus (40 cm wide × 40 cm long × 40 cm high) with a video recorder and analyzed by the ANY-Maze software (Stoeltling). Each animal (8-10 wks old) was individually tested and allowed to explore the arena for 5 minutes before the 10-minute testing session, as described previously (Gould et al., 2009). Wall climbs and rears, defined as when the subject raised its body only to be supported by its hind legs and tail, were quantified by an investigator who inspected the video and was blinded to the animal genotypes. The apparatus was cleaned with 70% ethanol and dried with paper towel after each animal was tested.

#### Rotarod Test

Mice were placed on the center rotating rod at a constant rotation of 4 repetitions per minute (rpm) which then accelerated within 30 seconds from 4 to 40 rpm. The maximum velocity (rpm) and the latency to fall (minutes) were recorded. Animals were tested three times with approximately 30 minutes between each testing interval, and the average of the three tests was reported.

#### Elevated Plus Maze

The maze consists of four plastic grey arms (39.5cm long × 5cm wide) connected perpendicularly; open arms alternate with a closed arm enclosed by walls (10cm tall). Metal legs attached to the end of each arm elevate the maze 50cm above ground (“ground”= a table). Testing began by placing a mouse in the center of the maze. The mouse was allowed to move freely while its body movements were recorded by ANY maze tracking software using an overhead camera for 10 minute sessions, and recording frequency at 10 positions/second. Duration of time spent in five different zones was measured, including the 4 arms and center of the maze. Zone entries and exits were scored, where entry was defined as a minimum of 70% of the animal’s body crossing into a zone, and not exiting was defined as a minimum of 30% of the animal remaining in the zone. Immobility was defined as 65% for a minimum of 2000ms. Freezing on threshold was set to 30 and freezing off threshold was set to 40 with a minimum duration of 250ms.

#### Marble burying

A mouse was placed in the middle of a new mouse cage filled with bedding 5cm high and 18 marbles arranged in 6 rows of 3, equidistantly spaced. The mouse was left inside for 30 minutes. The extent of marble burying was quantified as partially (<50%), mostly (>50%) and fully (100%), and the numbers of marbles buried were buried more than 50% in volume were called as buried.

### Surgery and EEG studies

Animals were anesthetized using isoflurane (4% in O_2_ for induction and 1.5-2% in O_2_ for maintenance) in a stereotaxic apparatus, where they were given 1 mL of 0.9% saline and 5 mg/kg Anafen for analgesia. For EEG recordings, 10- to 12-week-old mice were implanted with a two-tungsten microelectrode with polyimide tubing (one probe was placed into the frontal cortical regions, and one reference parietal electrode; wire diameter was 125 µm, length was 400 µm, MicroProbes, Gaithersburg, MD, USA). The electrodes were secured using cyanoacrylate (Krazy Glue) and Ortho-Jet acrylic (Central Dental Ltd., Scarborough, ON). Following surgery, the mice were given 1 mL of saline and 5 mg/kg of Baytril and monitored until they recovered completely from the anesthesia. All animals had a minimum post-operative recovery period of three days before recordings commenced.

#### EEG recordings

Mice were placed in warm plexiglass chambers (Harvard Apparatus, Holliston, MA) for a one-hour acclimation period, as described previously (Joshi et al., 2018). EEG activity was recorded in freely-moving animals for 90 minutes using a Grass amplifier and Grass software (Grass Instruments, Quincy, MA) and behaviors were monitored by a videorecorder. Mice were returned to their home cage, where they were then euthanized and brains harvested to determine electrode placement. The EEG recordings were obtained and analyzed from the bandwidth of 1 Hz low frequency filter (LFF) and 70 Hz high frequency filter (HFF) and with the 60 Hz notch filter ‘ON’. For Racine scale assessment with EEG confirmation, video episodes lasting 25 minutes during which animals exhibited extended immobility were selected. The number and time lengths of seizure episodes were quantified and assessed by a modified Racine scale: (1) stiffness and rigid posturing; (2) head nodding and tilting; (3) partial forelimb clonus; (4) whole body seizures; (5) motor convulsions, including falling, forelimb clonus and jumping (Kim et al., 2019; Racine, 1972).

### Magnetic Resonance Imaging

Mice were terminally anesthetized with a ketamine/xylazine mix and then transcardially perfused with 30 mL of 0.1 M PBS containing 10 U/mL heparin and 2 mM Prohance (gadolinium contrast agent; Bracco Diagnostics Inc.) followed by 30 mL of 4% paraformaldehyde (PFA) containing 2 mM Prohance. After perfusion, the mouse was decapitated and the skin, ears, eyes, and lower jaw were removed, leaving the brain within the skull, and incubated in 4% PFA/Prohance solution overnight at 4°C. The skull and brain were placed in a solution of PBS, 2 mM Prohance, and 0.02% sodium azide for 4 weeks before imaging. A 7.0-Tesla MRI scanner (Agilent Ince.) was used to acquire all images, and procedures have been described previously (Cahill et al., 2012; de Guzman et al., 2016). For the anatomical scan, a 40-cm inner bore diameter gradient was used (maximum gradient 30 G/cm) in conjunction with a custom-built solenoid coil array to image 16 samples in parallel (Lerch et al., 2011). To assess the volume differences throughout the brain a T2-weighted, 3-D fast spin-echo sequence, with a cylindrical acquisition of k-space, a TR of 350 ms, and TEs of 12 ms per echo for 6 echoes, field-of-view equaled to 20 × 20 × 25 mm3 and matrix size equaled to 504 × 504 × 630 was used. These parameters output an image with 0.040 mm isotropic voxels. The total imaging time for this sequence is 14 hours (Spencer Noakes et al., 2017).

Deformation-based morphometry (DBM) was used to analyze the volume differences throughout the brain. To visualize and compare any changes in the mouse brains, the images are linearly (6 followed by 12 parameter) and non-linearly registered together. Registrations were done with a combination of mni_autoreg tools and ANTS (advanced normalization tools) (Avants et al., 2008; Avants et al., 2011). Scans were resampled with the appropriate transform and averaged to create a population atlas representing the average anatomy of the study sample. The result of the registration is to have all images deformed into alignment with each other in an unbiased fashion. This allows for the analysis of the deformations, the goal being to model how the deformation fields relate to genotype (Lerch et al., 2008). The jacobian determinants of the deformation fields can then calculated as measures of volume at each voxel. Significant regional volume differences are calculated by warping a pre-existing classified MRI atlas onto the population atlas. The classified MRI atlas includes 182 different structures and incorporates several pre-existing atlases 182 different segmented structures encompassing cortical lobes, large white matter structures (i.e. corpus callosum), ventricles, cerebellum, brain stem, and olfactory bulbs (Dorr et al., 2008; Steadman et al., 2014). Multiple comparisons were controlled for using the false discovery rate (FDR) (Genovese et al., 2002).

### Histology

Mice were either anesthetized by hypothermia (P5 or younger), with an overdose of sodium pentobarbital (>P6 and older) or under isofluorane (4% in O_2_ for induction; 1.5-2% in O_2_ for maintenance), and transcardially perfused with cold phosphate buffered saline (PBS) followed by 4% PFA. Brains were post-fixed in 4% PFA overnight or for 2 hours at 4°C, and then rinsed and stored in PBS with 0.06% sodium azide. Postnatal brains (P14 and younger) were cryopreserved in 30% sucrose for 24 hours, embedded and frozen in OCT cryopreservation agent (Tissue-Tek), and coronal sections were collected at 20 µm using a Cryostat microtome. To prepared floating coronal sections from > P28 animals, brains were cryopreserved in 30% sucrose for 48-72 hours and sectioned at 35 µm. Sagittal cerebellar sections were prepared at 75 µm using a vibratome.

Primary and secondary antibodies were diluted in PBS in 10% normal donkey serum (NDS, Jackson Immunoresearch). Slidies with postnatal cryosections were incubated successively with blocking solution (0.3% Triton-X, 5% normal donkey serum in PBS) and primary antibodies (overnight at 4°C). Floating brain slices were rinsed with 0.5% PBS-Triton X and incubated in the blocking solution (10% NDS in PBS) for 1.5 hours at room temperature, then with primary antibodies for 48-72 hours at 4°C. After 3 × 10 minute washes in PBS, sections were incubated for 2 hours at room temperature with Alexa Fluor-conjugated secondary antibodies (Thermo Fisher Scientific or Jackson Immunoresearch). Nuclei were labeled with DAPI, or neuronal somata were stained with NeuroTrace 435/455 Blue Fluorescent Nissl (Thermo Fisher Scientific). Tissue was mounted and cover-slipped using Dako Mounting Medium (Agilent Technologies, Inc., CA, USA).

The primary antibodies used for this study are: goat anti-PV (1:2000, Swant, PVG-213); rabbit anti-SST (1:1500, Peninsula Laboratories International T-4103.0050); mouse anti-Reelin (1:2000, Millipore MAB5364), mouse anti-Gad67 (1:1000, Millipore, MAB5406), rabbit anti-VIP (1:1000, Immunostar 20077); rabbit anti-Calbindin (1:5000, Swant CB38); chicken anti-RFP (1:1500, Millipore AB3528); rabbit anti-RFP/DsRed (1:2000, Rockland 600-401-379); rabbit anti-cleaved Caspase-3 (1:300, Cell Signaling Technology 9661); rabbit anti-cleaved Caspase-9 (1:300, Cell Signaling Technology 9509); rabbit anti-FoxO3a (1:750, Cell Signaling Technologies 2497); mouse anti-synaptotagminII (1:300, Developmental Studies Hybridoma Bank, znp-1); rabbit anti-vGAT (1:2000, Synaptic Systems 131 003); mouse anti-Gephyrin (1:1000, Synaptic systems 147 011).

### Cell counting

For all quantifications, slides were coded and counts were performed with the examiner blind to the genotype.

#### Stereology

Sections from selected brain regions (Figure 2: somatosensory cortex, S1BF; globus pallidus, GP; basolateral amygdala, BLA; and primary visual cortex, V1) were imaged using a Leica SP8 confocal microscope and Fiji software (http://imagej.net/Fiji/Downloads) (Schindelin et al., 2012) was used to count cells. A section sampling fraction (ssf) for each region was determined, with the first section being randomly selected, allowing for an unbiased series of sections. After calculating the total area of the region, the optical dissector method was used for each region (grid size listed below) to select points within the region at 5x magnification. These points were then observed at a higher (63x) magnification to obtain optical z-stack images from each region. Cell counts were collected from each of these z-stack images at a thickness of 1.8 µm, for a total of ∼32 µm, and 1024 pixels by 1024 pixels. A guard zone of 3.6 µm at each superficial portion of the slice was selected to prevent bias due to super-saturation of cells or cutting artefacts. Each cell was identified and counted using an unbiased design-based stereological procedure. Total cell numbers for each region were estimated using the optical fractionator method (Mayhew and Gundersen, 1996). For the S1BF [Bregma +0.38 mm to −1.94 mm], the ssf = 1/6, the counting frame was set at 49.50 µm × 49.50 µm (PV) or 39.96 µm × 39.96 µm (NeuroTrace), and the grid size was 171.17 µm × 171.17 µm. For the GP [Bregma + 0.02 mm to −1.70 mm], the ssf = 1/4, the counting frame was set at 39.96 µm × 39.96 µm, and the grid size was 237.45 µm × 237.45 µm. For the BLA [Bregma - 0.58 to −2.06 mm], the ssf = 1/6, the counting frame was set at 49.50 µm × 49.50 µm, and the grid size was 158 µm × 158 µm. For the V1, the ssf = 1/7, the counting frame was set at 43.20 µm × 43.20µm, and the grid size was 132 µm × 132 µm.

#### Quantifications of cortical interneuron subpopulations

Composite images spanning the primary somatosensory cortex (SIBF) were acquired by stitching a grid of 9-12 confocal images with 20X objective and 0.8µm z-step size on a Leica SP8 (LasX Software, Leica) taking care that pairwise comparisons were acquired with the same acquisition settings. Cell counts for PV+, SST+, Reelin+, VIP+ classes were obtained within a specified region of interest (ROI, width: 1380 µm × height: spanning layers I-VI) and containing a maximum projection of 8 confocal z-stacks for Figure 3, or 8.0 um z-depth for other analyses. Cortical layers were determined with DAPI counterstaining and cell counts were subdivided into layers I-III, IV, and V-VI, using ImageJ and the cell counter plug-in. Areas of total and layer-specific ROIs were measured using ImageJ. Density was calculated as counts divided by the area (of the maximum projection).

For counts of Gad2-Cre; tdTomato-labeled interneurons through development, stitched images of barrel cortex were taken at 40x/1.4 NA. Using Imaris (BitPlane), images were segmented by cortical layers and ROIs were drawn based on DAPI staining to delineate cortical plate, layer V, layer VI in P2-P5 animals; or layer II/III, layer IV, layer V-VI in P8-P14 animals. The ROIs were extended in the z-axis to generate a 15 um thick volume, ensuring that this surface only encompassed imaged tissue (ie DAPI signal was present throughout the volume). This surface was used to mask the tdTomato channel, followed by spot detection on the tdTomato mask (estimated spot diameter of 8-12 um). Detected spots were manually validated to count only those spots with corresponding DAPI and tdTomato signals. For older time points, astrocytes were manually excluded based on morphology. To obtain a cell density value, the number of tdTomato+ spots was divided by the total volume of the masking surface for each layer, or summed across layers. A minimum of 4 ROIs per layer, per animal were analysed, and three mutant (Pcdhg^fcon3^ hmz; Gad2-Cre het; TdTomato het) and 3 control (Pcdhg het or wt; Gad2-Cre het; TdTomato het) genotypes. For apoptotic counts at P7, coronal sections spanning the somatosensory and visual cortical regions (Paxinos Developmental mouse brain atlas) were collected and simultaneously stained for cleaved Caspase-3 and Caspase-9. The tdTomato+/CC3-9+ signals were counted in the entire cortical sections.

#### Quantification of cerebellar molecular layer interneurons

For measurements of molecular layer thickness and Parvalbumin+ interneuron and Calbindin+ Purkinje cell number, 276 µm × 276 µm ROIs were sampled in folia 5 and 6. PV+ interneurons in the molecular layer were counted in three ROIs per cerebellar sagittal section, and the average was reported per section. Each counting frame consisted of one inclusion and one exclusion edge, and cells were counted if found entirely within the counting frame or overlapping with the inclusion edge but not the exclusion edge. For counts of Caspase-3+ apoptotic cells, images spanning the entire folia 5 were obtained and Z-stacks were collected with a 3µm step size. Caspase-3+ cells were counted within the molecular layer, and divided by the area of the maximum projection measured using the Fiji measurement tools.

### Image analysis

Immunofluorescence samples were imaged using a Leica SP8 confocal microscope equipped with 405, 488, 552, and 647nm lasers. Maximum projections were rendered from confocal stacks and contrast-enhanced using LasX (Leica), Photoshop (Adobe Systems) or Fiji (Schindelin et al., 2012). Modest non-linear enhancement was applied to some representative images. In some figures, inverted confocal images were presented to enhance clarity. Fluorescence intensity of FoxO3A immunostaining was analyzed using the surface detection algorithm in Imaris (BitPlane) to segment the tdTomato+ cells of interest in each image, followed by segmentation of DAPI-labeled nuclear surfaces. The FoxO3a+ channel was masked using both of these surfaces to segment cytosolic and nuclear compartments, and the ratio of nuclear to cytoplasmic fluorescence intensity was obtained to report the proportional FoxO3a subcellular localization.

### Single molecule FISH

HCR amplification based-single molecule FISH (smFISH) was performed following previous methods (Choi et al., 2010; Moffitt et al., 2016; Moffitt and Zhuang, 2016). P7 WT cortex was sectioned at 16 µm onto 12mm silane-treated coverslips. Primary probe hybridization targeting the Pcdhg constant region was performed at 2nM probe concentration in HCR hybridization buffer (Molecular Instruments, X) at 37 °C overnight. 4% polyacrylamide gels were cast for tissue clearing according to previous descriptions (Moffitt et al., 2016; Moffitt and Zhuang, 2016). Gels were allowed to incubate for 30 min at 4°C, prior to gelation at 37°C for two hours. Tissue clearing occurred overnight at 37°C in clearing digestion buffer (Moffitt et al., 2016; Moffitt and Zhuang, 2016). HCR amplification occurred at RT for 12 hours.

### Western Blot

Whole cortices from postnatal day 7 (P7) mice were dissected on ice, flash frozen in liquid nitrogen, and stored at −80°C until use. Cortices were homogenized in RIPA lysis buffer containing: 50 mM Tris pH 8, 150 mM NaCl, 2 mM EDTA, 0.5% sodium deoxycholate, 0.1% SDS, 1% NP-40 and 1× protease inhibitor cocktail (cOmplete, Roche), phosphatase inhibitor cocktails I and II (Millipore), and benzonase nuclease (10 KU, Novagen). Lysates were centrifuged at 12,000 rpm for 15 min and protein concentrations were determined by BCA assay (Pierce). Equal amounts of protein (11 µg) were run on a 10% SDS-PAGE gel and transferred to a polyvinylidene difluoride membrane (Millipore) overnight. Membranes were blocked for 1 h at room temperature in 5% BSA in TBST (20 mM Tris-HCl pH 7.5, 150 mM NaCl and 0.1% Tween20) and then incubated in primary antibody prepared in blocking buffer overnight at 4°C. The primary antibodies used for Western blot analyses were as follows: rabbit anti-P-Akt (Ser473, Cell Signaling, 4060, 1:1,500); rabbit anti-Akt antibody (Cell Signaling, 4691, 1:1,500); mouse β-actin (Cell Signaling Technology, 700, 1:10,000). Membranes were washed and incubated in appropriate secondary IRDye antibodies (1:10,000, LI-COR Biosciences, 926-32211 and 926-68070) in blocking buffer and visualized on a FC Odyssey Imaging System.

### Immunoprecipitation and western blotting from FACS-sorted cINs

To enrich the cIN population for pAKT western blot analysis, we used FACS to isolate TdTomato-expressing cells from *Pcdhg ^f^*; Gad2-cre; tdTomato transgenic mice at postnatal day 7. Harvesting of cortical cells was conducted as described for analytical flow cytometry experiments with the following modifications. After dissociation, 1ml of the cell suspension was layered onto 4ml of 22% Percoll (Sigma P1644)+PBS and spun at 400xg, 4°C, for 10min without centrifuge brakes engaged. The supernatant was removed, cells resuspended in 500µl 2% HI-FBS + PBS (ThermoFisher 12484028), and passed through a 40µm nylon mesh cell strainer (FisherScientific 352235). DAPI was added as a viability marker (ThermoFisher D1306; 5µM). Using a Beckman Coulter MoFlo Astrios VBYR cell sorter, an approximately equal number of TdTomato-expressing cells were collected from each genotype. Isolated cells were pelleted (400xg, 4°C) and resuspended in 100µl lysis buffer (ThermoFisher 87787) containing protease (ThermoFisher A32955) and phosphatase inhibitors (ThermoFisher 78442). Lysis cocktail was incubated on ice in the dark for 20min with intermittent vortexing. To remove debris, lysates were spun at 13,000xg, 4°C, for 10min. Supernatant was aliquoted, flash frozen and stored at −80°C prior to immunoprecipitation and western blotting. PanAKT was immunoprecipitated from the lysate using the rabbit anti-mouse Akt antibody (C67E7, Cell Signaling Technologies 4691) and the PierceTM Classic Magnetic IP/Co-IP kit (ThermoFisher 88804). The IP product was then run on a Western blot and probed using both the rabbit anti-mouse panAkt (1:1000) and Phosopho-Akt (Ser473, Cell Signaling Technologies 4060; 1:1000) antibodies, using Licor blocking buffer (LIC0927-70001). FACS isolated SVZ cells were included as positive controls. IgG raised in rabbit was used as a negative control (Cell Signaling 2729). To visualize bands the Licor IRDye 800CW (LIC-926-32211) and FC Odyssey imaging system were used. Band size was determined using the PageRuler Plus prestained ladder (Licor 26619).

### Analytical Flow

Population specific flow cytometric analysis of pAKT immunoreactivity in TdTomato-expressing cINs was conducted using *Pcdhg ^f^*; Gad2-cre; Rosa26lsl-tdTomato transgenic mice at postnatal day 7. Brains from two *Pcdhg ^f/f^* and control heterozygous *Pcdhg^+/f^* littermates were included in each experimental replicate. Cell dissociations were generated according to (Beaudoin et al., 2012) with the following modifications. Neocortical tissue was dissected in ice-cold Ca2+/Mg2+-free HBSS (ThermoFisher 14175095) containing: 10mM HEPES (Wisent 330-050-EL), 1mM Na-pyruvate (Wisent 600-0110-EL), 0.1% D-glucose (Sigma G7021). Tissue was cut into ∼2mm^2^ pieces and transferred to 4ml ice-cold Hibernate A, (ThermoFisher A1247501) containing 1x B27-TM supplement (ThermoFisher 17504044). Trituration was accomplished by passing tissue through a series of blunt tip stainless steel needles of descending gauge; 25, 22, 18 x4 each (McMaster-Carr Supply 75165A687, 75165A683, 75165A676). Cells were filtered through a 70µm nylon mesh cell strainer (FisherScientific 352340), pelleted by centrifugation at 400x *g*, and suspended in 5ml ice-cold PBS (Wisent 311-010-CL) + 4% PFA (Electron Microscopy Sciences 15710) for 20 minutes. Fixed cells were washed twice in 10ml PBS, permeabilized in 500µl 0.3% Triton-X 100 (Sigma T8787) PBS + 2% BSA for 5min, and incubated for 30min in the primary + secondary antibody solution: 2%BSA-PBS + Rabbit anti-mouse Phosopho-Akt (Ser473) antibody (Cell Signaling Technologies 4060; 1:50), Goat anti-Rabbit Alexa 488 secondary antibody (ThermoFisher A32731: 1:1000) and PureBlu Hoechst 33342 nuclear dye (Bio-Rad 135-1304). Cells were washed twice in PBS to remove excess antibody, resuspended in 500µl PBS, and passed through a 40µm nylon mesh cell strainer (FisherScientific 352235). Flow cytometry was conducted using a Becton-Dickinson LSRII-CFI, VBGR analyzer and data processed using FlowJo™ software v10.6.1. Approximately 10-15K TdTomato+ events were recorded in each flow analysis. Specificity controls included: unlabeled TdTomato+ cells and TdTomato+ cells + 2°-antibody/Hoechst 33342.

### Statistical analyses

Statistical analyses were performed in GraphPad Prism and SPSS. Quantifications of cell number were performed on mutant mice from multiple litters, with wild-type (Cre positive and Cre negative) or heterozygous littermates (Cre positive) serving as controls. The analyst was blinded to the genotypes until quantifications were complete. Means of two groups were compared using the two-tailed Student’s *t*-test on condition of equivalent variances determined by the ANOVA *F-*test, or with the Mann–Whitney non-parametric test. Means of multiple samples were compared using one-way ANOVA and posthoc Tukey test. Other statistical tests are specified in figure legends.

## ACKNOWLEDGEMENTS

This research was supported by an award from the Scottish Rite Charitable Foundation of Canada, a Canada Research Chair (Tier 2), a Sloan Fellowship in Neuroscience, and a CIHR Project grant to J.L.L; a University of Toronto Open Fellowship and a Restracomp PhD Student award (J.M.); an NSERC Undergraduate Research (A.A.); a Restracomp Postdoctoral Fellowship from the Hospital for Sick Children (A.D.S.); and the Province of Ontario Neurodevelopmental Disorders programme from the Ontario Brain Institute (J.P.L). We thank Giorgia Panichella for assistance on behavioral mouse analyses, and Dr. Krutika Joshi and Dr. Miguel Cortez (SickKids) for assistance for EEG recordings and analyses. We thank members of the Lefebvre laboratory for helpful comments on this manuscript.

## SUPPLEMENTARY FIGURES

**Figure 2 - supplement figure 1.**
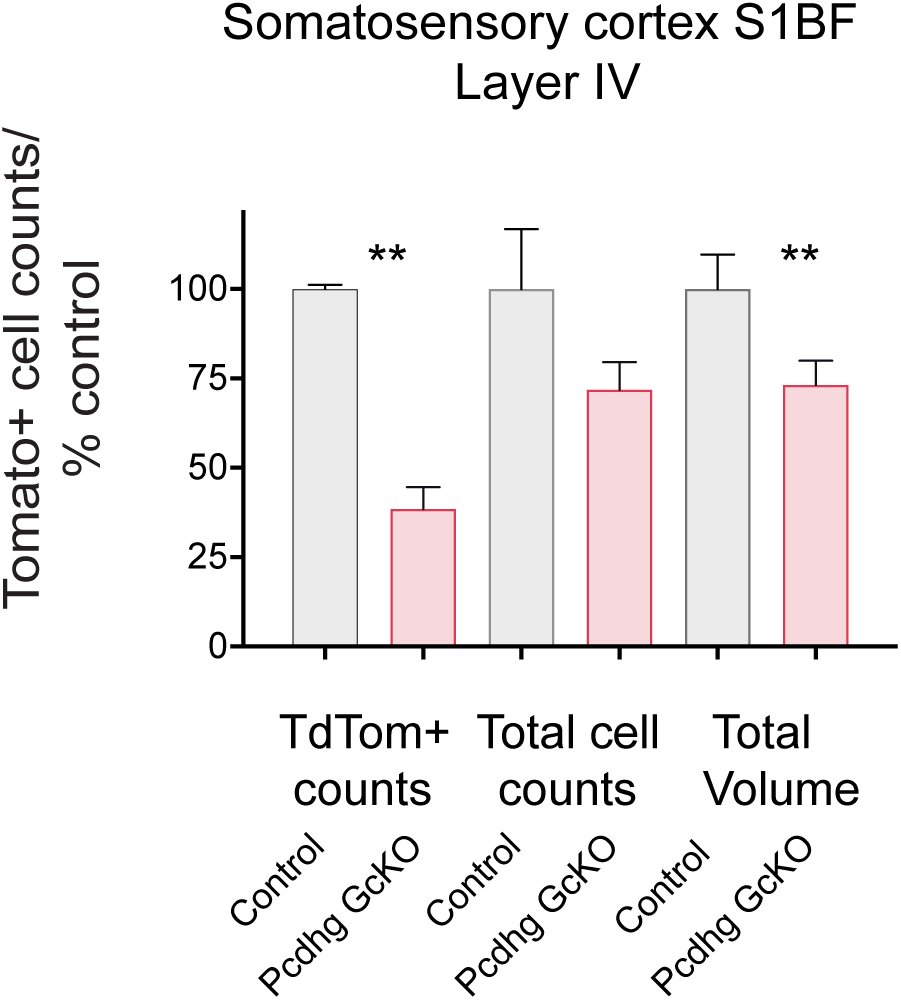
Numbers of Ai14-Tomato+ cINs are reduced in *Pcdhg GcKO* mutant. Stereological quantifications of Gad-Cre; TdTomato+ in Layer IV of somatosensory cortex (SIBF) of control and *Pcdhg GcKO* animals at P28. Data are normalized as percentage from control counts.

**Figure 5 - supplement figure 1.**
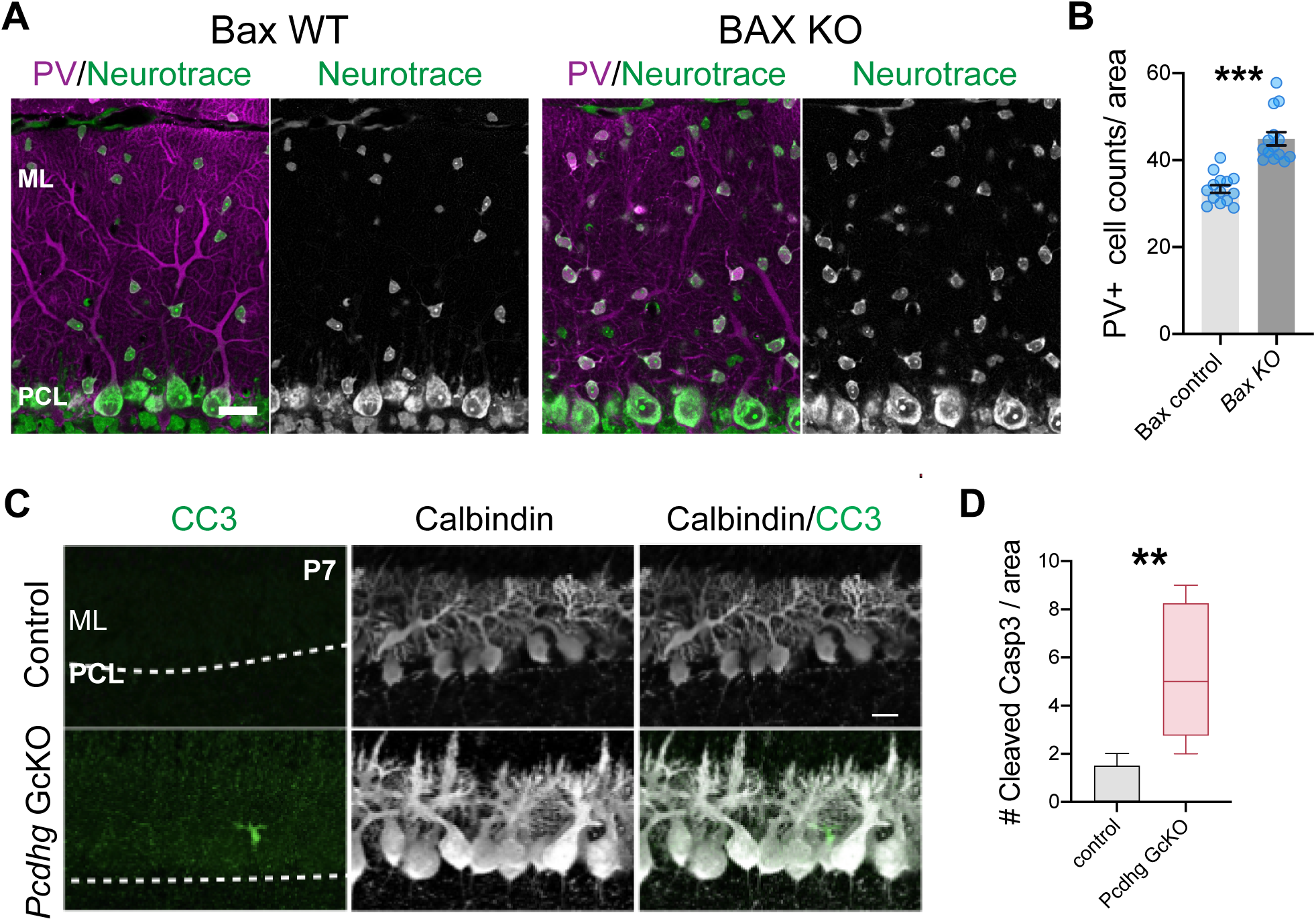
Cerebellar interneurons in the molecular layer undergo Bax-dependent developmental cell death and require the Pcdhgs for survival. **A.** Immunostaining of molecular layer interneurons (MLI) labeled by PV (magenta) and Neurotrace (NT, green) in cerebellar cortex of control and *BaxKO*. **B.** Quantifications of cerebellar PV/NT+ interneurons. Bars show mean +/− SEM and counts from 3 areas per section, 14 sections, 3 animals per genotype, *p* < 0.00001. Mann-Whitney U test. **C.** Immunostaining of cleaved caspase 3 (CC3, green) and Purkinje cell marker Calbindin (white) highlights an apoptotic interneuron (MLI) in the cerebellar molecular layer in *Pcdhg GcKO* at P7. **D.** Quantification of CC3 profiles in control and *Pcdhg GcKO* cerebellar cortex at P7, *p =* 0.001332, Mann-Whitney test. ** *p* < 0.01, *** *p* < 0.001. Scale bars: 20µm.

**Figure 5 - supplement figure 2.**
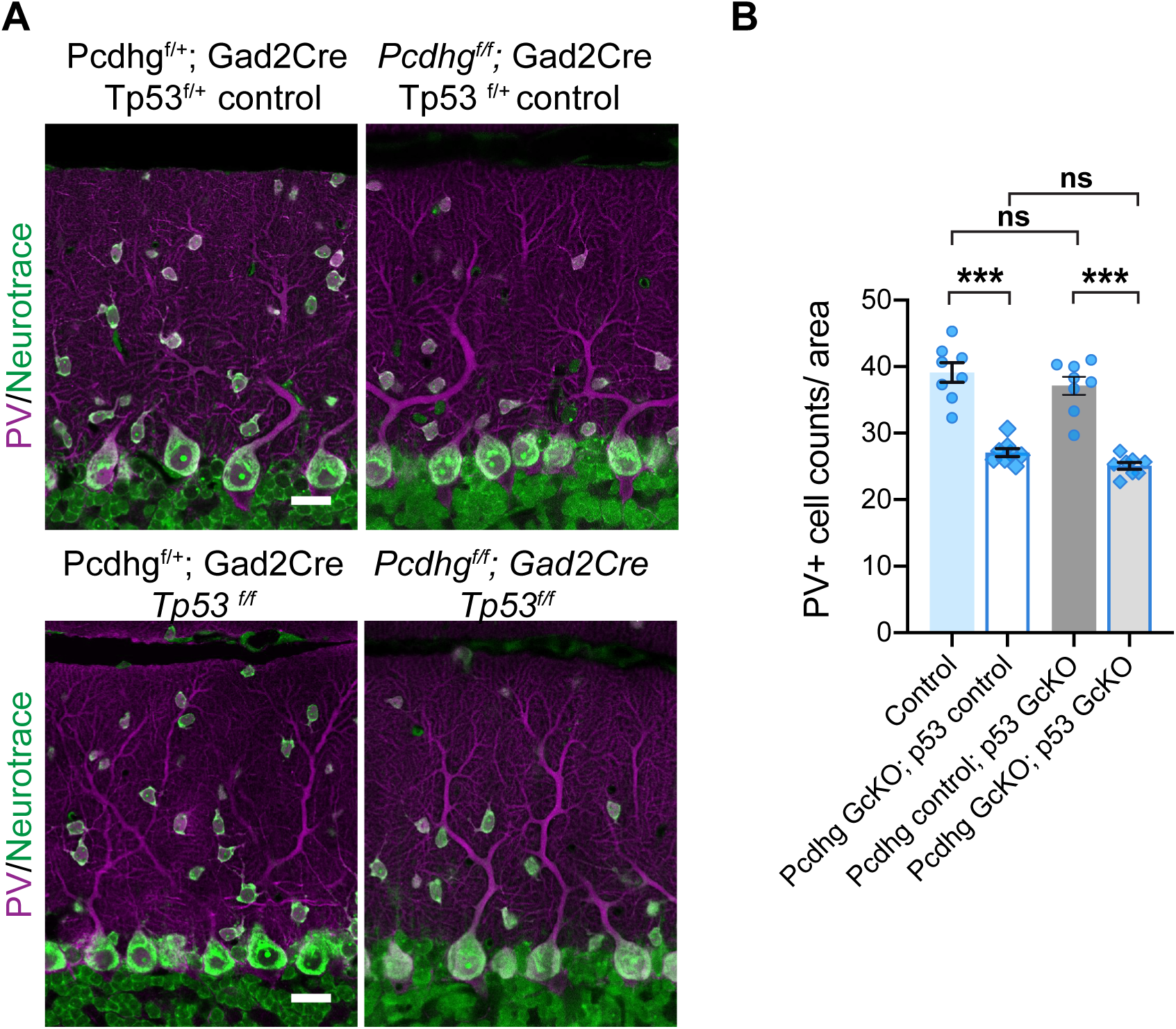
Cerebellar interneuron developmental cell death occurs through a p53-independent mechanism. **A.** Immunostaining of MLIs labeled by PV (magenta) and Neurotrace (NT, green) in cerebellar cortex of control, *Pcdhg GcKO; Tp53* control*, Pcdhg* het*; Tp53 GcKO*, and *Pcdhg GcKO; Tp53 GcKO* mutants. **B.** Quantifications of cerebellar PV/NT+ interneurons show no significance difference in *Tp53* homozygous mutants: Control vs. Pcdhg control; p53 GcKO, *p =* 0.36; Pcdhg GcKO; p53 control vs. Pcdhg GcKO; p53 GcKO, *p =* 0.36. Bars show mean +/− SEM and counts from 3 areas per section, 8 sections, 2 animals per genotype, *p* < 0.00001. Mann-Whitney U test. *** *p* < 0.001. Scale bars: 20µm in A.

**Figure 6 – supplement figure 1.**
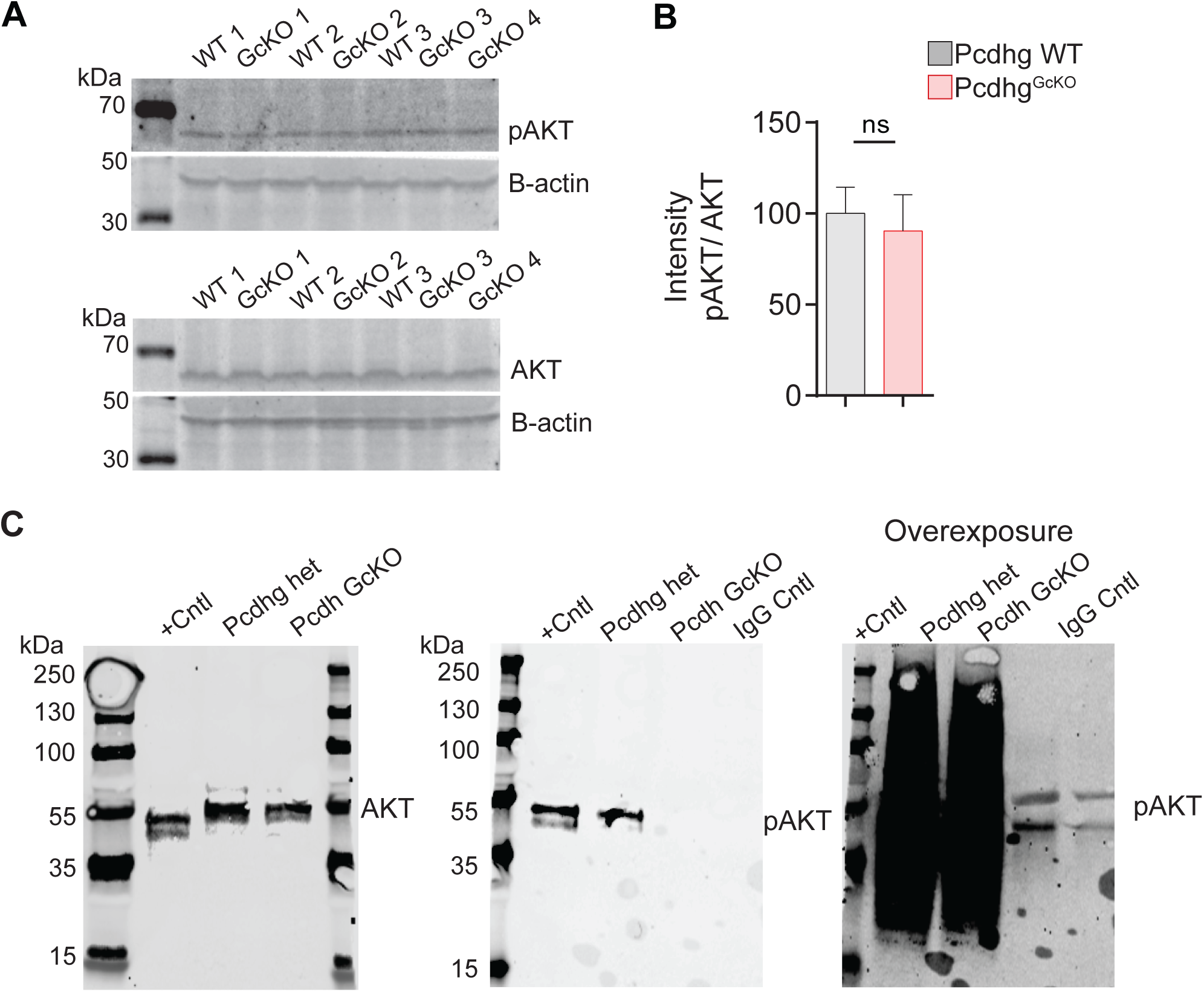
Detection of reduced phosphorylated AKT levels in *Pcdhg GcKO* cINs isolated by FACS. A. Immunoblots of AKT (top) and phosphorylated AKT Ser473 (bottom) of wild-type (WT) and *Pcdhg GcKO* P7 cortical lysates. Beta-actin loading control was used in both blots. B. Quantifications and relative intensity of pAKT and total AKT. Bars show pAKT/AKT ratio with SEM from technical replicates using WT, N = 6; *GcKO*, N = 7 animals; p = 0.34, unpaired t test with Welch’s correction. C. Immunoblots of AKT (left) and pAKT (middle; right = overexposure) following AKT-mediated immunoprecipitation of Gad-Cre; TdTomato+ cortical cells isolated by Fluorescence-activated cell sorted (FACS) from P7 Pcdhg het and *Pcdhg GcKO* cortical lysates. Positive control (+Cntl) is a validated sample. N = 2 animals per genotype. Equal numbers of Tdtomato+ cells from FACS were assayed.

**Figure 7 – supplement Figure 1.**
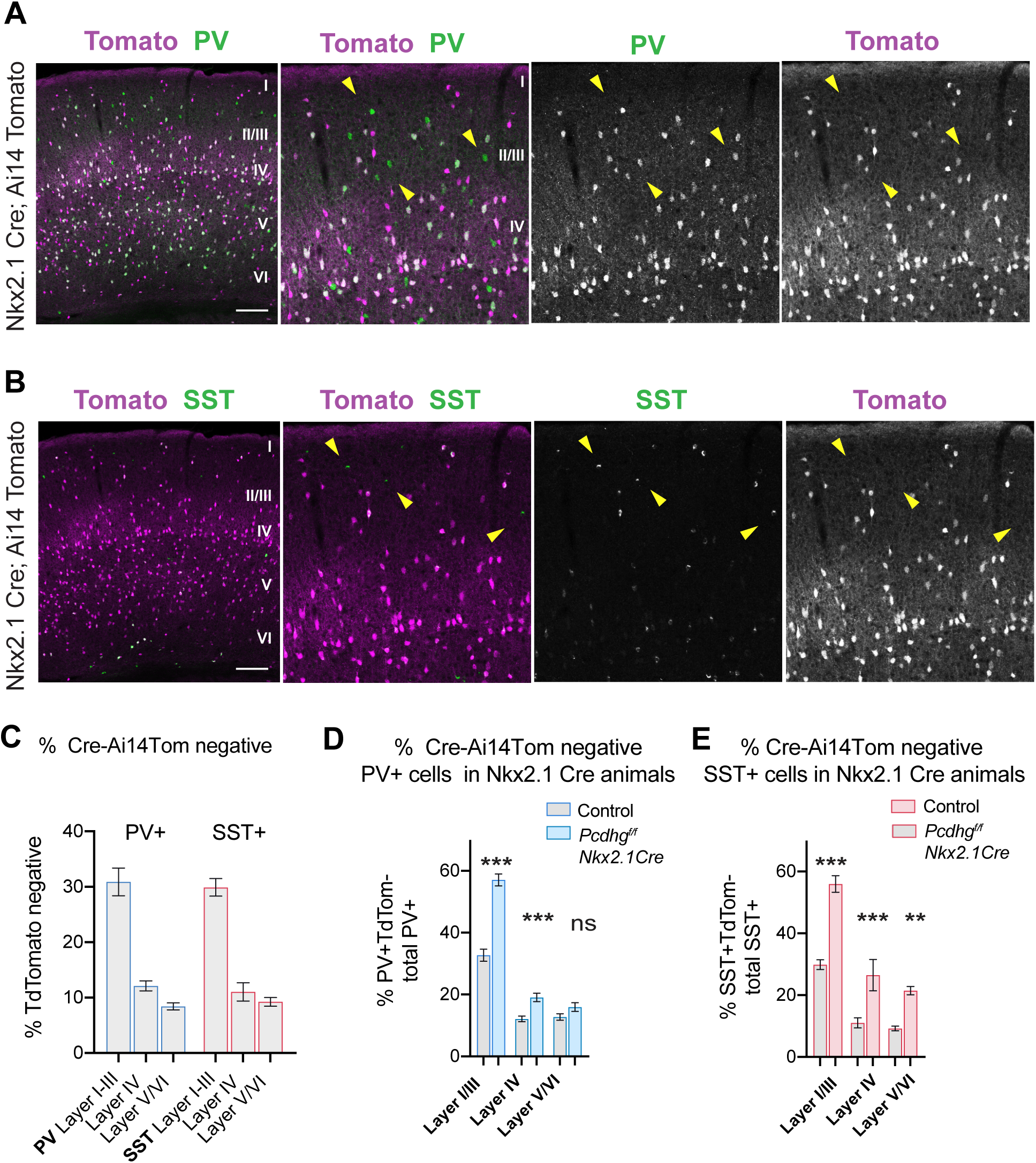
Mosaic recombination activity of Nkx2.1-Cre to test for Pcdhg autonomy. **A.** Immunofluorescence staining of SSC cortex of Nkx2.1-Cre; Ai14-LSL-TdTomato animals at P28 showing Cre+;TdTomato+ (magenta) and PV+ (green). Note overlapping PV+/Tomato+ and non-overlapping PV+/Tomato-negative (yellow arrows) cells. **B.** Immunofluorescence staining of SSC cortex of Nkx2.1-Cre; Ai14-LSL-TdTomato animals showing Cre+;TdTomato+ (magenta) and SST+ (green). Note overlapping and non-overlapping SST+/Tomato-negative (yellow arrows). **C.** Quantifications of PV+/Tomato-negative (blue) and SST+/Tomato-negative cells (red) in control Nkx2.1-Cre; Ai14-LSL-TdTomato SSC cortex at P28, represented as percentage of total PV+ or SST+ quantified in the same region. **D.** Quantifications of PV+/Tomato-negative cells [Cre negative thus Pcdhg WT] in SSC cortical layers in control and Pcdhg*^f/f^*; Nkx2.1-Cre; Ai14-LSL-TdTomato animals, represented as percentage of total PV+ cells in region of interest. PV+ I/III, *p* <0.00001; IV, p = 0.0005, V/VI, ns, not significant. Mann-Whitney U test. **E.** Quantifications of SST+/Tomato-negative cells [Cre negative thus Pcdhg WT] in SSC cortical layers in control and Pcdhg*^f/f^*; Nkx2.1-Cre; Ai14-LSL-TdTomato animals, represented as percentage of total SST+ cells in region of interest. SST+ I/III, *p* <0.00001; IV, p = 0.003, V/VI, p<0.00001. Mann-Whitney U test. Bars show mean +/− SEM from 12-15 sections, f4 animals per genotype. * *p* < 0.05; ** *p* < 0.01; *** *p* < 0.001. Scale bars: 200µm.

